# HiFi long-read metagenomics reshapes genome-resolved interpretations of activated sludge microbiome

**DOI:** 10.64898/2026.11.18.767492

**Authors:** Min Joon Song, Taeho Yun, Sojung Yoon, Ju Yong Lee, Ameet J Pinto, Sukhwan Yoon

**Author notes:** Corresponding author: Sukhwan Yoon.

## Abstract

Genome-resolved metagenomics is widely used to characterize activated sludge microbiomes and infer the microbial processes underpinning system performance. However, the reliability of metabolic inferences drawn from fragmented short-read (SR) genomes remains insufficiently examined. Using (near-)complete genomes reconstructed from HiFi long-read (LR) sequencing as references, we traced genome recovery through SR assembly and post-assembly processing. Assembly fragmentation and subsequent sequence loss during post-assembly processing compromised the recovery of high-quality genomes despite adequate sequencing depth and breadth of coverage. This incomplete recovery biased inferences regarding denitrification modularity, underestimating complete denitrifier populations and distorting the inferred balance between N_2_O producers and reducers. Poor recovery of nitrifier and Ca. *Accumulibacter* genomes, exacerbated by population microdiversity and structurally complex genomic regions, precluded reliable inferences regarding nitrification and phosphorus accumulation. Our findings caution against uncritical interpretation of SR-based microbiome-function linkages and highlight the value of incorporating HiFi LR sequencing into genome-resolved metagenomic analyses.

## Introduction

Wastewater treatment plants (WWTPs) are essential infrastructure for urban water sustainability, treating wastewater generated by households and industries worldwide^1^. Activated sludge (AS), a central biological component of modern WWTPs, harbors diverse and complex microbial communities, which underpin essential nutrient removal processes^2^. Spatial separation or temporal alternation of conditions differing in oxygen availability enables the coordinated operation of nitrification, denitrification and enhanced biological phosphorus removal^3^. Accordingly, advancing our understanding of the physiology and ecology of the microbial populations responsible for these processes is essential for improving nutrient removal performance and mitigating unintended consequences, such as N_2_O emissions^4^.

Over the past decade, metagenomics has been widely used for interrogating the structure and functional potential of AS microbiomes. Most published studies of AS metagenomes have relied on Illumina sequencing platforms, which, despite their advantages in cost-effectiveness, throughput, and base-calling accuracy, typically generate reads of 300 base pairs (bp) or shorter. Genome-scale resolution remains limited by assembly fragmentation, which compromises the accuracy and sensitivity of subsequent genome binning^5–7^. Activated sludge communities present additional complexities that challenge metagenomic assembly and binning, including high microdiversity^8,9^, uneven sequencing depth^8^, conserved operon structures^10^, repetitive and exogenous sequences^11,12^, and pervasive mobile genetic elements^13,14^, which collectively compromise assembly contiguity and undermine binning accuracy. Recent large-scale genome-resolved analyses of AS microbiomes have revealed extensive genomic diversity while highlighting how these compounding challenges hinder genome recovery^2^. Nevertheless, the extent to which incomplete genome recovery distorts inferences regarding key nitrogen- and phosphorus-cycling pathways and thereby biases interpretations of the microbial processes underpinning treatment performance remains insufficiently examined.

Long-read (LR) metagenomics, with recent technological advances, offers a promising means to improve genome-resolved metagenomic analyses. By applying circular consensus sequencing (CCS), single-molecule real-time (SMRT) sequencing can now achieve low error rates (<0.1%) with long read lengths (up to ∼25 kbp)^15^, enabling high-throughput recovery of (near-)complete genomes that can meet all high-quality draft criteria defined by the Minimal Information about a Metagenome-Assembled Genome (MIMAG) standards^16^. Advances in long-read metagenomic assembly tools, now supporting HiFi LR sequencing (e.g., hifiasm-meta, metaMDBG, and metaFlye), have mitigated assembly challenges posed by structurally complex genomic regions, thereby improving assembly contiguity and binning fidelity^8,17,18^. Moreover, highly contiguous assemblies enable the adoption of completeness-aware strategies that reduce erroneous binning assignments^19,20^. Accordingly, several studies applying HiFi LR shotgun metagenomics to complex microbial communities, e.g., human gut microbiomes, have demonstrated superior recovery of MAGs in terms of both yield and assembly quality compared with SR metagenomics^19,21^. Additionally, (near-)complete LR MAGs provide valuable references for systematically tracing genomic information loss during SR assembly and post-assembly processing.

Such genomic information loss may have particularly important consequences for interpreting the microbial processes underpinning biological nutrient removal. Nitrifier genomes are notoriously difficult to reconstruct^22,23^, whereas incomplete recovery of denitrification and phosphorus accumulation genes in SR MAGs may lead to erroneous interpretations of their metabolic capabilities. Nevertheless, the potential biases introduced by incomplete genome recovery are often inadequately considered when genome-resolved metagenomic or metatranscriptomic analyses are used to infer microbial contributions to nutrient removal performance or N_2_O emissions^4,24^. Here, using (near-)complete genomes reconstructed from HiFi LR sequencing as references, we (i) systematically traced genome recovery through SR assembly and post-assembly processing to identify the major causes of incomplete genome recovery and (ii) evaluated how incomplete genome recovery biases inferences regarding the metabolic capabilities of key microbial populations involved in biological nutrient removal. Our findings highlight how incomplete genome recovery can distort functional interpretations of AS microbiomes and underscore the value of HiFi LR metagenomics for improving the reliability of genome-resolved analyses.

## Materials and methods

Additional methodological details, parameter settings, and validation procedures are provided in the Supplementary Methods.

### Sample collection and DNA extraction

Activated sludge samples were collected on December 18, 2024 from Daejeon Municipal WWTP (36°23’5’’N 127°24’2’’E; South Korea), which operates with a typical anaerobic-anoxic-oxic (A2O) configuration. The operating parameters and performance of the WWTP can be found in our previous publication^25^. One-liter wastewater samples were grab-sampled from the oxic segments of four parallel trains and immediately transported to the laboratory in ice. Genomic DNA was extracted from 100 mL of each sample using DNeasy PowerSoil Pro Kit (QIAGEN GmbH, Hilden, Germany).

### Library preparation and sequencing

For PacBio HiFi long-read sequencing, 4 µg of gDNA, quantified with Quant-iT PicoGreen™ dsDNA assay kit (Invitrogen, CA) and dissolved in 10 µL of buffer, was fragmented using a Megaruptor 3 (Diagenode, NJ). The quality of the fragmented DNA was subsequently assessed using Femto Pulse system (Agilent, CA). Libraries were prepared with SMRTbell prep kit 3.0 and purified using AMPure PB beads (Pacific Biosciences, CA). Short-read shotgun sequencing libraries were prepared according to TruSeq Nano DNA Library Prep Reference Guide (15041110D). Both LR and SR sequencing were performed by Macrogen Inc. (South Korea), on PacBio Revio (24.8±0.9 Gbp; CCS-processed HiFi reads) and Illumina NovaSeq X (17.0±1.3 Gbp; raw short reads) platforms, respectively.

### Construction of metagenome-assembled genomes (MAGs)

PacBio circular consensus sequences were generated using the CCS module in SMRT Link v25.3 (Pacific Biosciences). HiFi reads from all four samples were pooled and assembled using three assemblers in parallel: *metaMDBG* v1.1, *hifiasm-meta* v0.1 and *metaFlye* v2.9.5 (--pacbio-hifi --meta)^8,17,18^. These HiFi reads and contigs were processed using HiFi-MAG-Pipeline v2.0 workflow (provided by PacBio; available at https://github.com/PacificBiosciences/pb-metagenomics-tools/blob/master/docs/Tutorial-HiFi-MAG-Pipeline.md). Resulting MAGs were dereplicated at the species level using *<u>dRep</u>* v3.5.0 (-sa 0.95 -nc 0.65). Genome quality estimates using *CheckM2* v1.1.0 were used in place of the default *CheckM* estimates^26,27^.

The NovaSeq-generated short-read sequences were processed using a workflow analogous to that applied to the LR dataset. The raw paired-end reads were trimmed using *fastp* v0.24.0 (-l 100 -q 25 −2)^28^. The paired-end sequences from four samples were pooled and assembled using *metaSPAdes* v4.1.0^29^. The quality-trimmed paired-end reads were mapped to the contigs using *Bowtie2* v2.5.4 and sorted using *SAMtools* v1.21^30,31^. The contigs were binned using *SemiBin2* v2.2.0 and *MetaBAT2* v2.17 and processed with *DAS Tool* v1.1.7 and *dRep* (with the same flags as for LR sequencing)^32–34^.

Sequences classified as viral or plasmid by *geNomad* v1.11.0 (score ≥0.9, ≥1 marker gene, and marker_enrichment ≥5), except for proviral sequences, were removed from both MAG catalogues^35^. Bins identified as chimeric by *GUNC* v1.0.6 were also removed^36^. Genome completeness and contamination were estimated using *CheckM2* v1.1.0, and low-quality bins (completeness<50% or contamination≥10%) were discarded, in accordance with the MIMAG standards^16^. Ribosomal RNA genes (5S, 16S, and 23S rRNA genes) in the MAG bins were predicted using *Infernal* v1.1.5 against Rfam release 15.0 with the option “--cut_ga --rfam --nohmmonly --oskip"^37,38^. Transfer RNAs (tRNAs) were identified using *tRNAScan-SE* v2.0.12, applying the bacterial or archaeal model based on the taxonomy assigned by the *classify_wf* workflow in *GTDB-Tk* v2.6.1 against the GTDB release 226^39^. Genome circularity and structural integrity were assessed using an expanded read-to-assembly validation framework for HiFi LR assemblies, as detailed in Supplementary Methods^40^. The resulting MAGs were classified into quality categories using criteria adapted from the MIMAG standards (Supplementary Table 1)^16^.

### Genome characterization and annotation

Relative abundances of the MAGs were estimated by mapping quality-trimmed SR sequences to the MAG catalogue using *CoverM* v0.6.1 (options: -p bwa-mem2 --min-read-percent-identity 95 --min-read-aligned-percent 65)^41^. Ribosomal RNA genes were extracted using the *getfasta* module in *BEDTools* v2.31.1 and searched against *MiDAS* v5.3 database using *BLASTN* v2.17.0^42,43^. Repeated regions were identified using BLASTN self-alignments with predefined length and nucleotide-identity thresholds, as detailed in Supplementary Methods. Taxonomy was assigned using *classify_wf* workflow in *GTDB-Tk* v2.6.1 based on GTDB release 226^44^.

Coding sequences were predicted using *Prodigal* v2.6.3, run in single-genome mode^45^. Complete genes were defined as coding sequences possessing both start and stop codons^46^. Genes were annotated using *KofamScan* v1.3.0 against KOfam database (released 2025-12-01)^47^, and only significant hits were retained for downstream analysis. Additional functional annotation was performed to complement KO assignment using *InterProScan* v5.75-106.0^48^, *bakta_proteins* module in *Bakta* v1.11.2 (database v6.0)^49^, *DIAMOND BLASTP* v2.1.24 searches against NCBI clusteredNR protein database (2026-02-12) and the Swiss-Prot database (2026-02-13)^50^. NorB family proteins (K04561), which were not reliably identified by these approaches, were identified by screening predicted protein sequences with *hmmscan* in *HMMER* v3.4 against a publicly available set of NorB HMMs (https://github.com/ranjani-m/HCO)^51^. Metabolic pathways were inferred using *kegg-pathways-completeness-tool* based on the presence of KEGG Orthologs^52^. Only KEGG modules with 100% completion were considered present.

### Reference-guided assessment of short-read genome recovery

The LR MAG catalogue comprising 447 genomes served as the reference for competitive SR read recruitment and contig assignment. Read coverage breadth, mean read depth, and depth heterogeneity were quantified from primary alignments. Within-population microdiversity was quantified using *inStrain* v1.10.0^53^. High-confidence SR contigs assigned to each LR MAG were used to quantify assembly breadth (fraction of the LR MAG covered) and contiguity (N50). Gaps between SR contig alignments and candidate misassemblies identified by *QUAST* were assessed using HiFi read support^54^. Assigned SR contigs were traced through initial binning, consolidation, dereplication and the final SR MAG catalogue construction. Read support, microdiversity and the fractions occupied by repeats, insertion sequences, paralogous regions, and proviruses were compared between poorly (<50% assembly breadth) and near-completely (≥95%) recovered 1-kbp windows.

### Comparison of conspecific genome pairs between platforms

Conspecific LR and SR genome pairs were identified to compare genome- and gene-scale recovery between sequencing platforms and to trace the loss of genomic information during assembly and post-assembly processing of SR data. The 182 (near-)complete LR MAGs were compared against the SR catalogue using *skani* v0.3.1 dist -- slow^55^. Genome pairs with average nucleotide identity (ANI) ≥95% and aligned fractions (AF) ≥65% in both directions were considered conspecific. Insertion sequences were predicted using *ISEScan* v1.7.3. Pairwise comparisons of orthogroup presence and copy number were performed using *OrthoFinder* v2.5.5^56^. Additionally, 39 nitrification-related loci from eleven (near-)complete nitrifier LR MAGs were evaluated for potential homologous-copy collapse.

### Phylogenomic analysis

For each (near-)complete MAG, bacterial SCGs were extracted using the *identify* module in *GTDB-Tk* v2.6.1. Amino acid sequences corresponding to these SCGs were aligned separately for each gene using *MAFFT* v7.525 with the --auto option^57^. The resulting alignments were trimmed using *trimAl* v1.5.0 with the heuristic mode (-automated1)^58^. The individual SCG alignments were concatenated into a phylogenomic supermatrix using *create_concatenation_matrix* command in *PhyKIT* v2.0.2^59^. A maximum-likelihood phylogeny was inferred from the concatenated alignment using *IQ-TREE* v3.1.4, with the best-fitting substitution model selected using *ModelFinder* and branch support assessed with 1,000 ultrafast bootstrap and 1,000 SH-like approximate likelihood ratio test replicates^60^. The resulting phylogeny was visualized using R package *ggtree* v3.20^61^.

### Statistical analysis

Statistical analyses were performed in R v4.5.2 using the built-in *stats* package and *correlation* package v0.8.0^62,63^. Paired Wilcoxon signed-rank tests were applied to conspecific genome comparisons, within-genome window classes and nitrifier read-depth comparisons. Pairwise associations among the analysed variables were assessed using Spearman’s rank correlations. Partial Spearman correlations were calculated using rank-transformed variables to account for covariation among predictors. Statistical significance was set at 0.05, with *P* values adjusted using the Benjamini-Hochberg procedure.

## Results and discussion

### Long-read metagenomics enables recovery of complete prokaryotic genomes

The LR and SR workflows yielded 811 and 167 draft MAGs, respectively, which were reduced to 448 and 148 non-redundant MAGs after species-level dereplication. Inspection of within-species variation (>99% ANI and >65% AF) in the pre-dereplicated genomes identified 44 additional strain-level variants uniquely resolved by LR metagenomics, demonstrating its capacity to resolve microdiversity beyond the species level. After exclusion of seven chimeric SR genomes and one LR genome during manual curation, the final datasets comprised 447 LR and 141 SR genomes (Fig. 1a; Supplementary Data 1). Genome-wide read recruitment supported all recovered MAGs, with minimum breadths of 87.8% and 99.98% and minimum mean depths of 1.8× and 6.3× for the LR and SR MAGs, respectively (98.4% breadth and 5.0× depth for the 182 (near-)complete LR MAGs; Fig. 1b). The low overall coverage breadth for LR MAGs likely resulted from competitive read mapping, in which reads shared by closely related MAGs were assigned to only one MAG. The proportions of quality-trimmed SR reads mapped to recovered genomes were substantially higher for the LR dataset (37.4±1.4%; 38.8±0.8% without dereplication) than for SR dataset (16.3±1.0%), indicating an improved genome recovery by LR metagenomics (Fig. 1c). For both the LR MAG catalogue (Spearman ρ = 0.94; *P*<0.05) and the SR MAG catalogue (ρ = 0.90; *P*<0.05), relative abundance estimates based on LR- and SR-read mappings were strongly correlated (Fig. 1c).

**Fig. 1:**
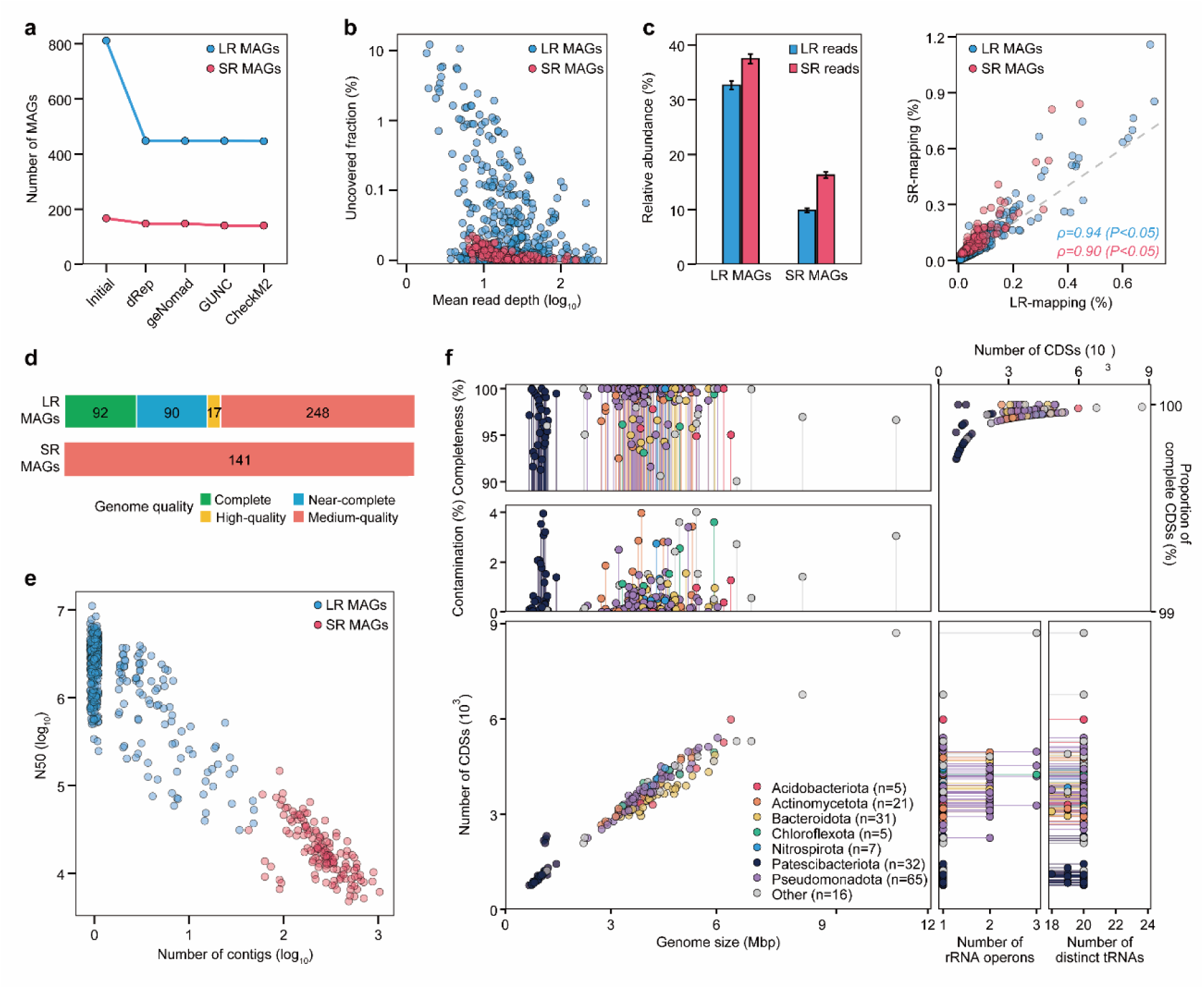
Recovery and quality statistics of LR and SR MAGs. **a,** Changes in MAG counts through post-assembly processing stages**. b,** Mean read depth and the fraction of genomic positions with zero read depth in each sample, to which PacBio HiFi and Illumina reads were mapped, respectively. Read depth and breadth were calculated from primary and supplementary alignments. **c,** Cumulative relative abundance of the organisms represented by LR and SR MAGs, estimated by LR and SR read recruitment (left), and Spearman’s correlation between relative abundances of the MAGs estimated from two read types (right). Error bars indicate standard deviations. **d,** Distribution of LR and SR MAGs across MIMAG quality categories. **e,** Relationship between the numbers of contigs and N50 of individual MAGs. **f,** Genomic characteristics of complete and near-complete LR MAGs, including completeness and contamination calculated with *CheckM2* v1.1.0 (upper left), proportion of protein-coding sequences with start and stop codons (upper right), and numbers of complete rRNA operons and unique tRNAs (bottom).

**Extended Data Fig. 1.**
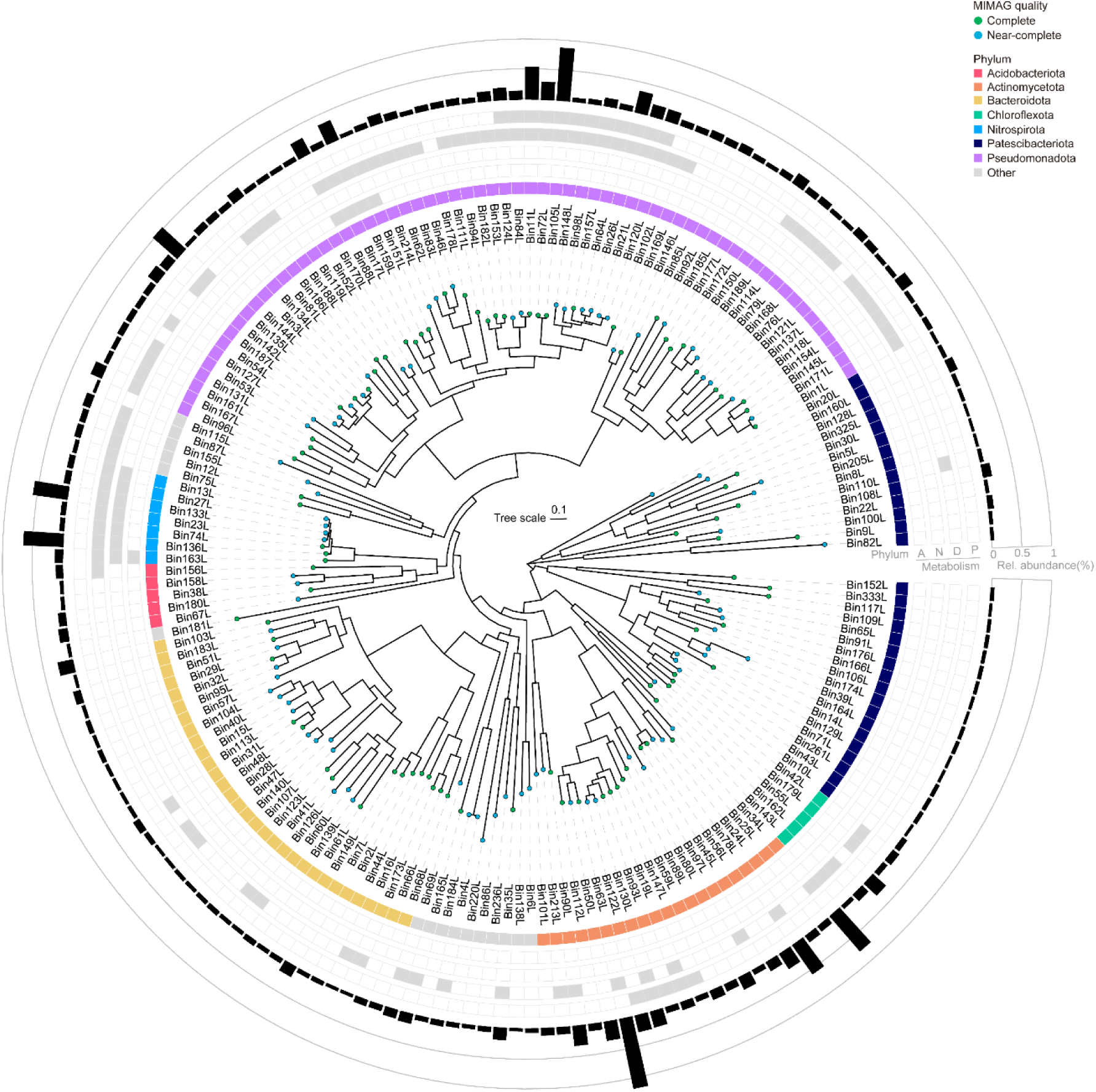
Phylogenomic tree of 182 (near-)complete MAGs reconstructed with HiFi LR metagenomic sequencing. The colors of the tree tip points indicate genome quality (complete or near-complete). The innermost annotation ring indicates phylum-level taxonomic affiliation. The four subsequent concentric rings indicate the presence (grey) or absence (white) of genetic potential for ammonia oxidation (A), nitrite oxidation (N), canonical denitrification (D), and enhanced biological phosphorus removal (P). The outermost ring indicates the mean relative abundance of each MAG in the four activated sludge samples, as estimated by SR mapping.

Complete and near-complete genomes were recovered exclusively from the LR dataset. Of the qualified 447 LR MAGs, 92 were classified as complete (20.6%) and 90 as near-complete (20.1%) genomes, whereas all SR MAGs were classified as medium-quality (Fig. 1d). This difference was primarily attributable to substantially greater assembly contiguity (Supplementary Note 1): 350 LR MAGs comprised a single contig, whereas all SR MAGs remained fragmented, with an average N50 of 22.7 kbp (Fig. 1e). The LR MAGs showed average genome completeness and contamination values of 81.2±18.2% and 1.4±2.2%, respectively. At least one full-length copy of each of the 5S, 16S, and 23S rRNA genes was recovered in 365 LR MAGs, and the LR MAGs encoded tRNAs corresponding to 17.6±3.1 of the 20 standard amino acids. Genome integrity assessment detected no excessive repeats, false duplications, or false closures in the LR MAG catalogue, whereas phantom sequences occurred in 10 MAGs, including six that otherwise met the criteria for (near-)complete genomes (Supplementary Note 2; Supplementary Data 2). All 92 complete genomes showed assembler-reported circularity, which was further validated by HiFi read mapping. Seven additional MAGs (four affiliated with the genus *Promineifilum*, one with the genus *Azoamicus*, and one each with the phyla Cyanobacteriota and Patescibacteriota) showed read-supported circularity without evidence of false closure but were classified as medium-quality based on other genome-quality criteria (Supplementary Note 3).

The complete and near-complete genomes recovered by LR metagenomics represented extensive genomic diversity within AS microbiomes (Fig. 1f; Extended Data Fig. 1). These genomes all represented distinct species (ANI<95% or AF <65%) and collectively accounted for 18.7±1.2% of the community. Genome sizes ranged from 0.7 to 11.1 Mbp with 753 to 8,708 protein-coding genes, indicating the pronounced genome streamlining or expansion among specific lineages (Fig. 1f). These extremes included streamlined Patescibacteriota genomes (1.0±0.2 Mbp) and an exceptionally large, GC-rich Myxococcota genome (11.1 Mbp; Supplementary Note 4 and Supplementary Fig. 1). Nearly all of 605,692 predicted CDSs (99.9±0.1 %) were recovered with both start and stop codons.

Taxonomic classification based on GTDB indicated that the genomes spanned 16 bacterial phyla, with *Pseudomonadota* (n=65), *Patescibacteriota* (n=32), *Bacteroidota* (n=31), *Actinomycetota* (n=21), and *Nitrospirota* (n=7) being the most represented. A substantial fraction lacked close taxonomic representatives in GTDB, with 13 genomes lacking genus-level matches and 127 lacking species-level matches. Even among the 55 genomes with species-level matches in GTDB, 53 were previously represented by multi-contig assemblies (217±273 contigs; N50=261±532 kbp; Supplementary Fig. 2; Supplementary Data 3).

### Cross-platform genome comparison reveals the sources of incomplete short-read genome recovery

To examine the extent of incomplete genome recovery by SR metagenomics, we compared 56 conspecific SR MAGs with their (near-)complete LR counterparts at the genome and gene level (99.8±0.2% ANI; reciprocal AFs of 81.5±8.3% and 95.3±5.2% for LR-to-SR and SR-to-LR MAGs comparisons, respectively; Supplementary Data 4). Repetitive regions, mobile elements, paralogous loci, and CRISPR arrays were disproportionately underrepresented in SR MAGs (Supplementary Note 5; Supplementary Fig. 3). Copy-number discordance was observed in 7.6±3.9% of orthogroups, of which 71.8% were either detected only in LR counterparts or present at higher copy numbers in LR counterparts (Supplementary Note 6; Supplementary Fig. 4; Supplementary Data 5). These differences occurred despite a mean *CheckM2*-estimated completeness of 90.2±6.9% for the SR MAGs, showing that high estimated genome completeness can obscure substantial loss of genomic content, including protein-coding sequences potentially relevant to genome-based functional inference.

Comparison of the conspecific genome pairs showed incomplete SR assembly despite nearly complete coverage of LR MAGs by SR reads (99.8±0.2% at ≥1×; mean depth, 24.7±25.6×; Supplementary Note 7; Supplementary Data 6)^29,64^. SR contigs competitively aligned to the 447 LR MAGs collectively covered only 84.0±10.6% of LR MAG length across the 56 conspecific pairs (Fig. 2a). At least two HiFi reads spanned 95.9% of the 10,291 gaps between regions aligned to SR contigs, suggesting that longer reads are needed to resolve regions prone to SR assembly fragmentation. HiFi read support indicated that QUAST may have overestimated structural misassemblies, as 43 of the 51 candidate misassembly junctions were supported by HiFi reads. Post-assembly processing resulted in an additional loss of 7.9±4.3% of LR MAG length, with bin consolidation as the main culprit, yielding a final recovery of 76.1±12.1% in the paired SR MAGs. Of the SR contigs assigned to each LR MAG, 9.2±5.0% was not retained in the final SR MAG catalogue, and 0.5±0.7% was placed in non-conspecific SR MAGs.

**Fig. 2:**
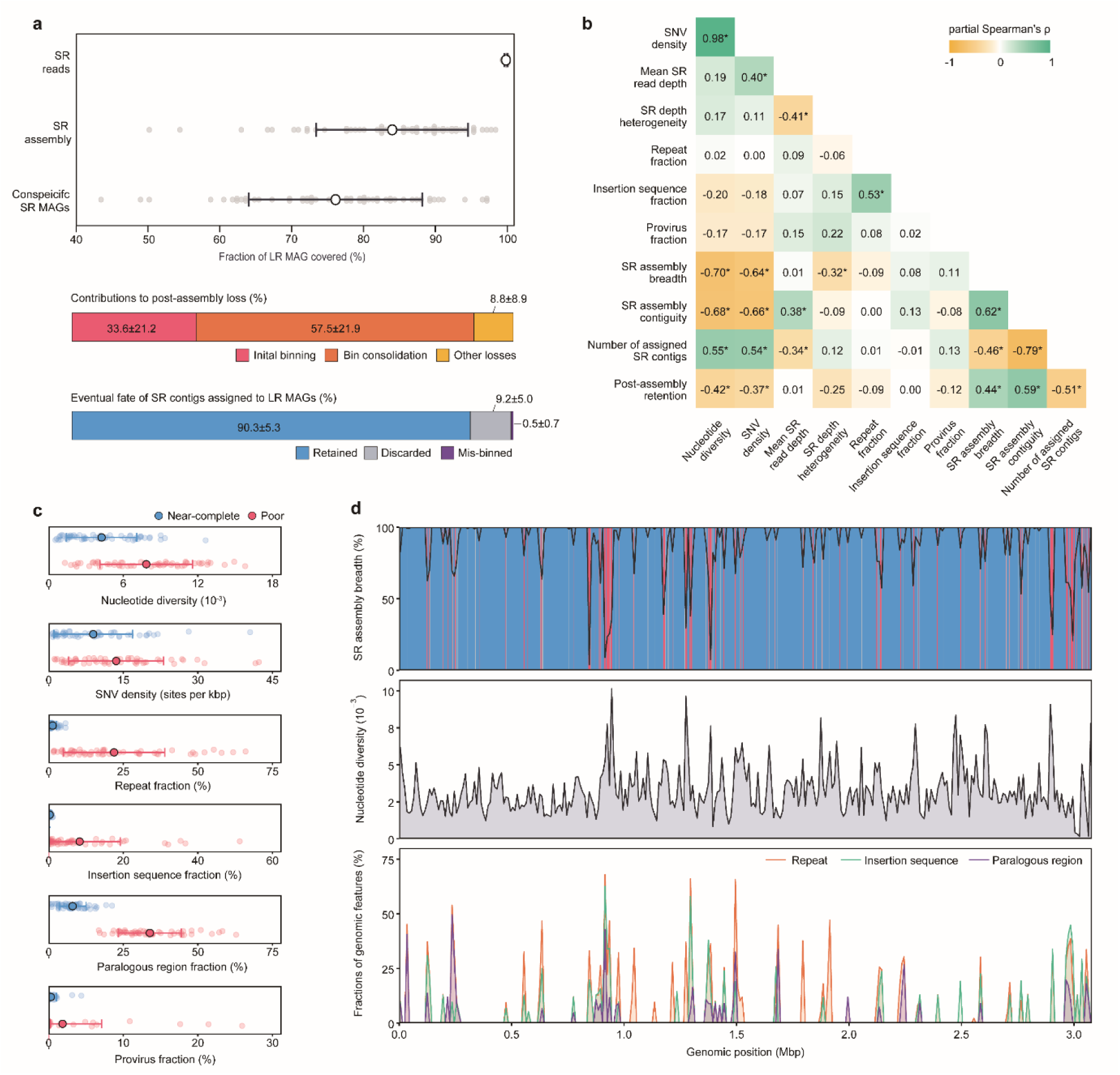
Genome-wide and local factors underlying incomplete short-read genome recovery. **a,** Fractions of LR MAG sequences covered by SR reads, assemblies and conspecific SR MAGs (top). Point-range plots show means±s.d, and jittered points represent individual MAGs. Contributions of individual post-assembly processing steps to sequence loss (middle) and the eventual fate of SR contigs assigned to LR MAGs (bottom) are presented as stacked bar graphs. **b,** Partial Spearman correlations among genome metrics. Asterisks indicate significant correlations (*P*<0.05). **c,** Comparative analyses of 1-kbp windows in LR MAGs with near-complete (≥95%) and poor (<50%) recovery by SR assembly. Jittered points represent individual 1-kbp windows. **d,** Genome-wide profiles of SR assembly breadth (top), nucleotide diversity (middle), and frequencies of genomic features (repeats, insertion sequences, paralogous regions, and provirus; bottom) along the near-complete genome of *Methylophosphatis* (bin124L). Blue and red shading indicate local regions of near-complete (≥95%) and poor (<50%) SR assembly recovery, respectively. Recovery classes were defined in 1-kbp windows, whereas line profiles represent values smoothed over 10-kb windows.

Genome-wide nucleotide diversity and SNV density consistently showed significant negative correlations with both SR assembly breadth (partial Spearman’s ρ=-0.70 to −0.64) and contiguity (ρ=- 0.68 to −0.66; all *P*<0.05; Fig. 2b; Supplementary Note 8). Poorly recovered 1-kbp windows showed higher SNV density (13.6±9.5 versus 9.0±7.9 sites per kbp; *P*<0.05) than near-completely (≥95%) recovered windows and were also enriched in repeats, insertion sequences, paralogous regions, and proviruses (Fig. 2c; all *P*<0.05), as illustrated by the *Methylophosphatis* genome pair (Fig. 2d).

### Accurate depiction of denitrification modularity requires LR metagenomics

Long-read metagenomics demonstrated clear evidence that denitrification, i.e., the reduction of NO ^-^ or NO ^-^ to gaseous nitrogen products NO, N_2_O, and N_2_, in AS is dominated by partial denitrifiers that collectively constitute a modular pathway. A total of 302 LR MAGs, including 71 complete and 68 near-complete genomes harbored at least one of the genes responsible for a denitrification step, *narG*, *napA*, *nirK*, *nirS*, *norB*, and *nosZ*, with 31.1±1.2% of the quality-trimmed short reads mapped onto these MAGs (Fig. 3a; Supplementary Data 7). A substantial proportion of these microorganisms (135 MAGs; 10.0±0.8% relative abundance) lacked the genetic capacity for NO ^-^-to-NO reduction, a defining criterion of canonical denitrifiers, suggesting that microorganisms traditionally not classified as denitrifiers may nonetheless contribute substantially to denitrification at the community level.

**Fig. 3:**
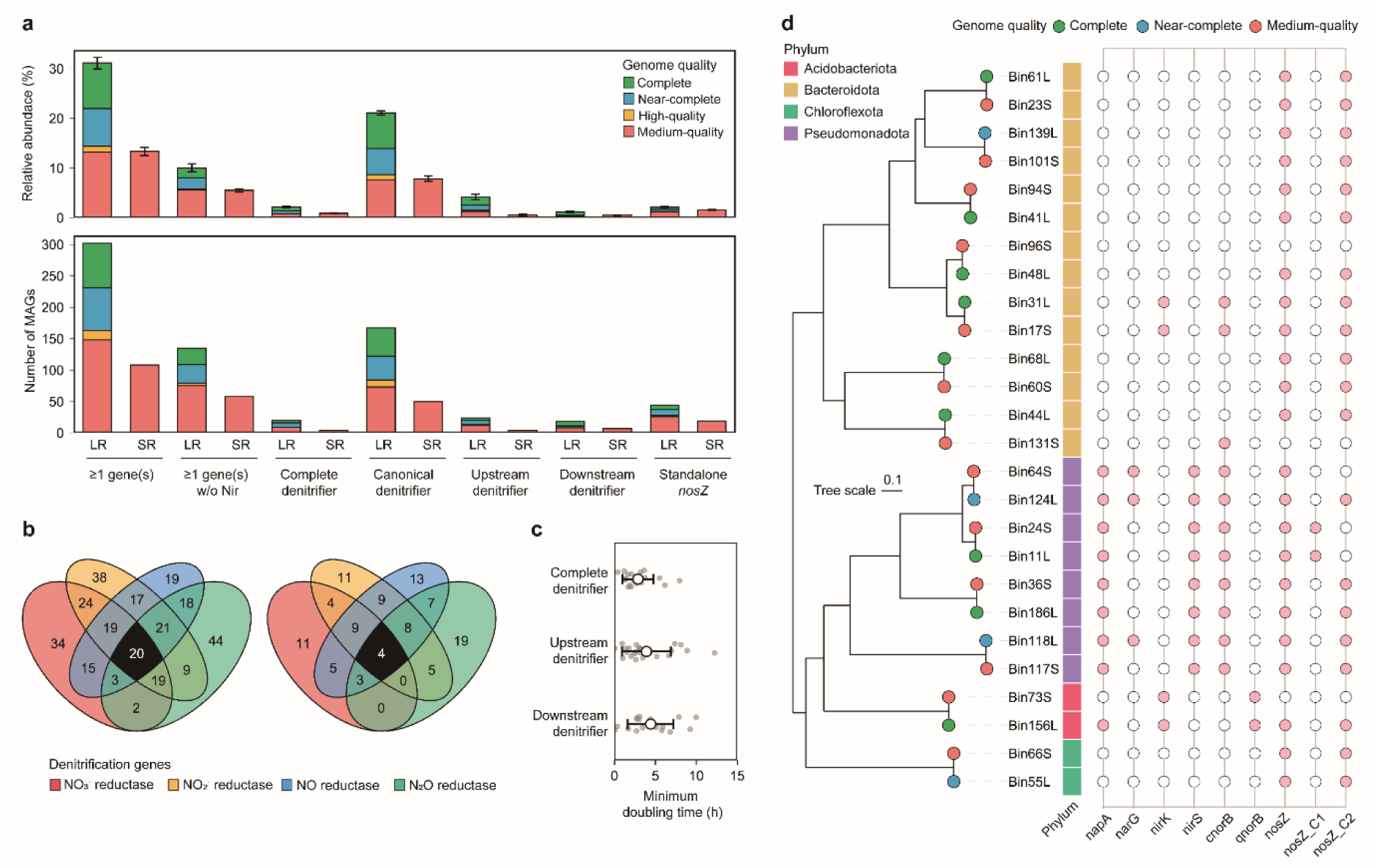
Comparison of LR versus SR metagenomics in characterizing denitrifying population. **a,** The cumulative relative abundances (top) and counts (bottom) of LR and SR MAGs encoding each denitrification step, stratified by their quality. Upstream and downstream denitrifiers were defined as MAGs encoding genes for only the two upstream (NO ^-^ and NO ^-^ reduction) and two downstream (NO and N_2_O reduction) denitrification steps, respectively. Note that the categories are not mutually exclusive. **b,** Venn diagram showing the distribution of LR (left) and SR (right) MAGs according to their genetic potential for the four denitrification steps. Numbers indicate MAG counts belonging to each category. **c,** Minimum doubling times of complete, upstream and downstream denitrifiers predicted using *gRodon2* v2.5.2. Points indicate individual MAGs and the error bars indicate standard deviations of the MAG-level estimates within each category. **d,** Phylogenomic tree of LR MAGs encoding the NO_2_^-^-to-N_2_ pathway or standalone *nosZ* and their conspecific SR counterparts. Filled pink and open circles indicate gene presence and absence in the MAGs, respectively. nosZ_C1 and nosZ_C2 denote clades I and II, respectively.

**Extended Data Fig. 2.**
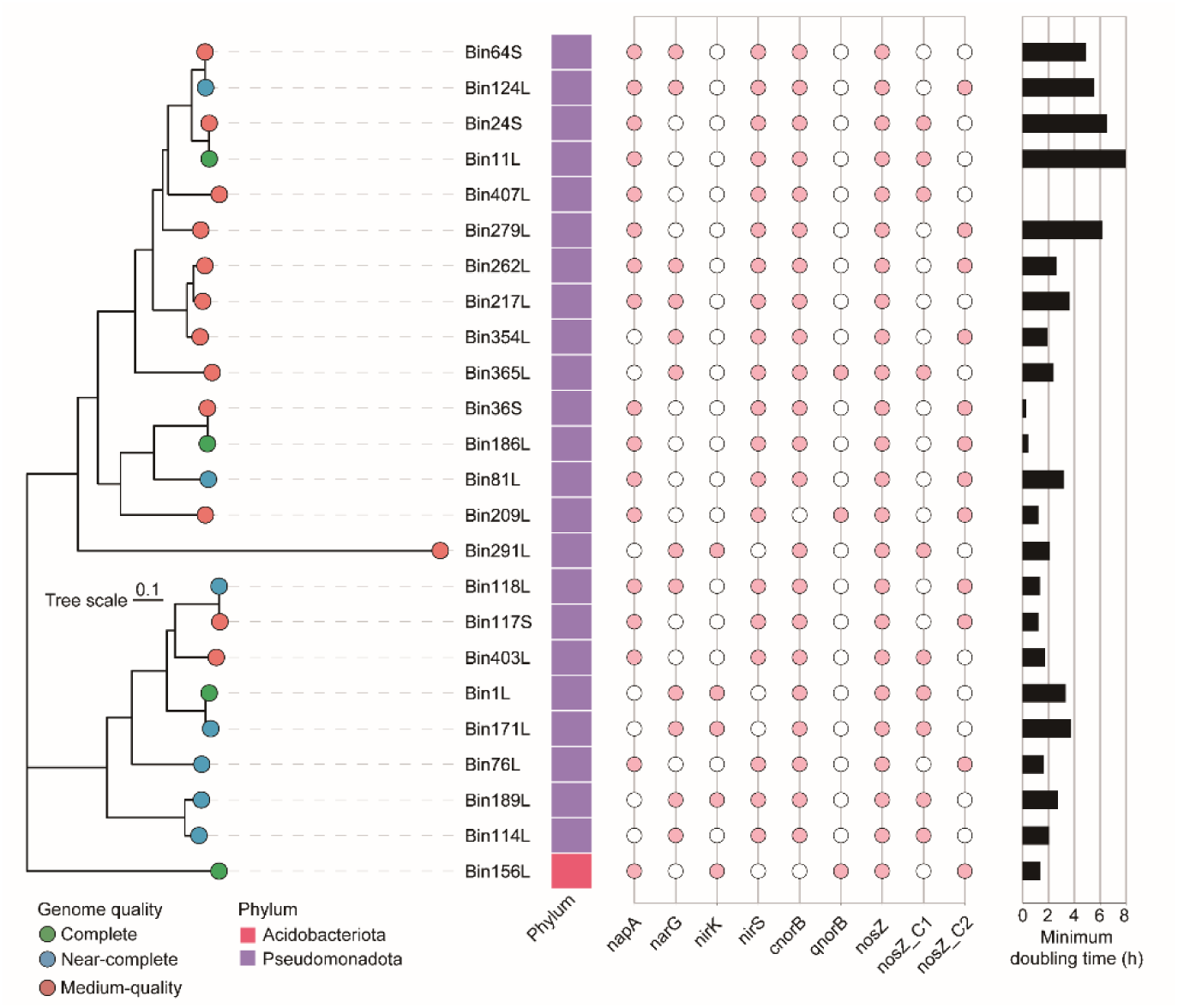
Phylogenomic diversity of complete denitrifiers. Phylogenomic tree of LR and SR MAGs classified as complete denitrifiers. Filled pink and open circles indicate gene presence and absence, respectively. *nosZ* clades I and II are denoted by nosZ_C1 and nosZ_C2, respectively. Horizontal bars show minimum doubling times predicted using *gRodon2*.

**Extended Data Fig. 3.**
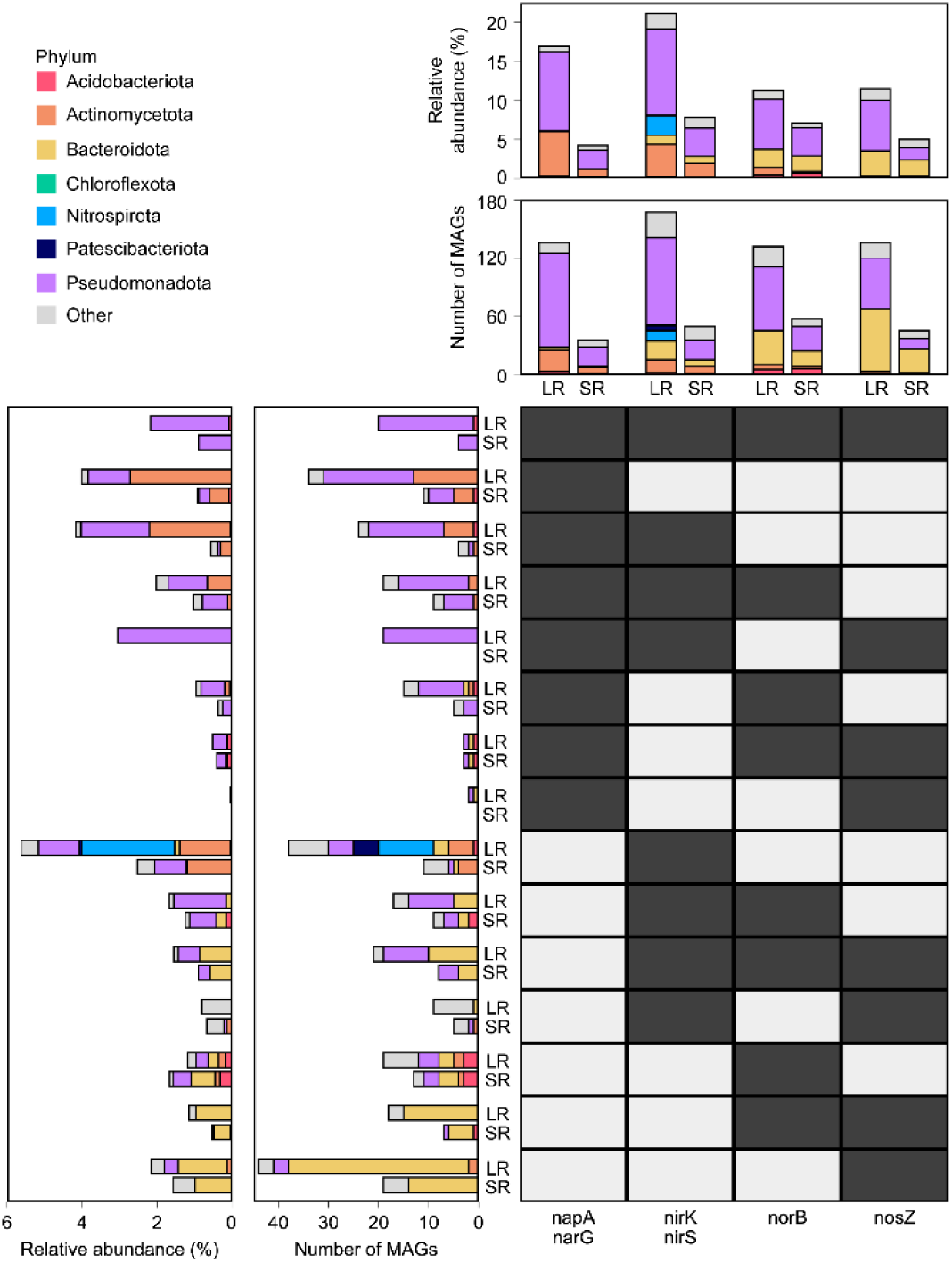
Comparison of denitrifier gene repertoire recovery between LR and SR metagenomics. Dark and light cells in the matrix indicate the presence and absence, respectively, of genes associated with each denitrification step. The vertical stacked bars above the matrix show cumulative relative abundances (top) and numbers of MAGs (bottom) with genomic potential for each denitrification step. The horizontal stacked bars to the left of the matrix show the corresponding metrics for individual combinations of denitrification steps.

Only a small subset of the LR MAGs (4 complete, 7 near-complete, and 9 medium-quality MAGs with 2.2±0.2% of total short reads mapped) encoded a complete gene set for NO ^-^-to-N denitrification (Fig. 3b and Extended Data Fig. 2). The complete denitrifiers were dominated by Pseudomonadota (7 Alphaproteobacteria and 12 Gammaproteobacteria). The only complete denitrifier genome outside this phylum was affiliated with the order Vicinamibacterales within the phylum Acidobacteriota.

However, their low relative abundance alone does not establish that denitrification flux is predominantly mediated by modular pathways, because metabolic activity cannot be inferred from genome abundance alone. Consistent with the caveat, growth rate predictions using *gRodon2* indicated that complete denitrifiers exhibit shorter predicted minimum doubling times (2.9±1.9 h) than both upstream (3.9±3.0 h) and downstream (4.4±2.8 h) partial denitrifiers, although these differences were not statistically significant (*P*>0.05; Fig. 3c).

Beyond addressing the fundamental limitation that the true absence of genes encoding specific denitrification steps cannot be reliably inferred from the fragmented incomplete MAGs that dominate SR datasets^12,65^, long-read metagenomics markedly improved the resolution of modular denitrification by enabling a substantially more comprehensive recovery of genomes encoding *nirK* and *nirS*. The LR workflow recovered 167 MAGs with 21.1±0.4% relative abundance, substantially exceeding the recovery achieved by the SR workflow (50 MAGs with 7.9±0.5% relative abundance). This broader recovery included 16 *nirK*-harboring MAGs affiliated with Nitrospirota (n=11) and Patescibacteriota (n=5), which together constituted 2.6±0.4% of the community and were recovered exclusively by the LR metagenomes (Extended Data Fig. 3).

One of the most conspicuous differences between the LR and SR MAG sets, with important implications for greenhouse gas emissions from WWTPs, was the recovery of *nosZ*-harboring MAGs, either as part of a NO ^-^-to-N denitrification pathway or with *nosZ* as a standalone denitrification gene. Often, the quantitative comparison between the organisms with standalone *nosZ* versus those with upstream denitrification genes encoding N_2_O-producing nitrite and nitric oxide reductases is used as a predictor of N_2_O emission potential of a microbiome^66,67^. Of the 41 LR MAGs encoding

NO ^-^-to-N pathway, collectively representing 3.7±0.2% of the community, one of six conspecific SR counterparts (bin73S; 0.12±0.03%) lacked *nosZ* and would therefore be misclassified as an N_2_O producer (Fig. 3d). Among seven SR MAGs conspecific to the *nosZ*-only LR MAGs, two lacked *nosZ*, one of which (bin131S) encoded a spurious *cnorB* gene, similarly contributing to an erroneous metric. These results indicate that an SR MAG-based estimate of N_2_O production–consumption potential can be biased by both false-negative gene loss and false-positive gene assignment. In summary, LR metagenomics is required for reliable quantitative comparison of partial versus complete denitrifier populations and a robust genetic prediction of ecosystem-scale N_2_O production and consumption potential.

### Long-read metagenomics reveals hidden features of nitrifier genomes

Eleven complete and near-complete genomes affiliated with nitrifiers collectively accounted for 2.2±0.3 % of the total community, comprising four *Nitrosomonas*, six canonical *Nitrospira*, and one comammox *Nitrospira* (Fig. 4a). All *Nitrosomonas* genomes encoded three *amoCAB* operons and three NH_2_OH ubiquinone redox modules (HURM; *haoAB*-*cycAB*) with the exception of bin88L, which harbored two *amoCAB* operons. All canonical NOB encoded one to three *nxrAB* clusters. The sole comammox *Nitrospira* genome (bin163L, complete) encoded a single *amoCAB* operon, a single HURM module, and two *nxrAB* clusters (Supplementary Note 9). At least one copy of *nirK* was predicted in all these nitrifier genomes, although its physiological function outside AOB is yet unknown.

**Fig. 4:**
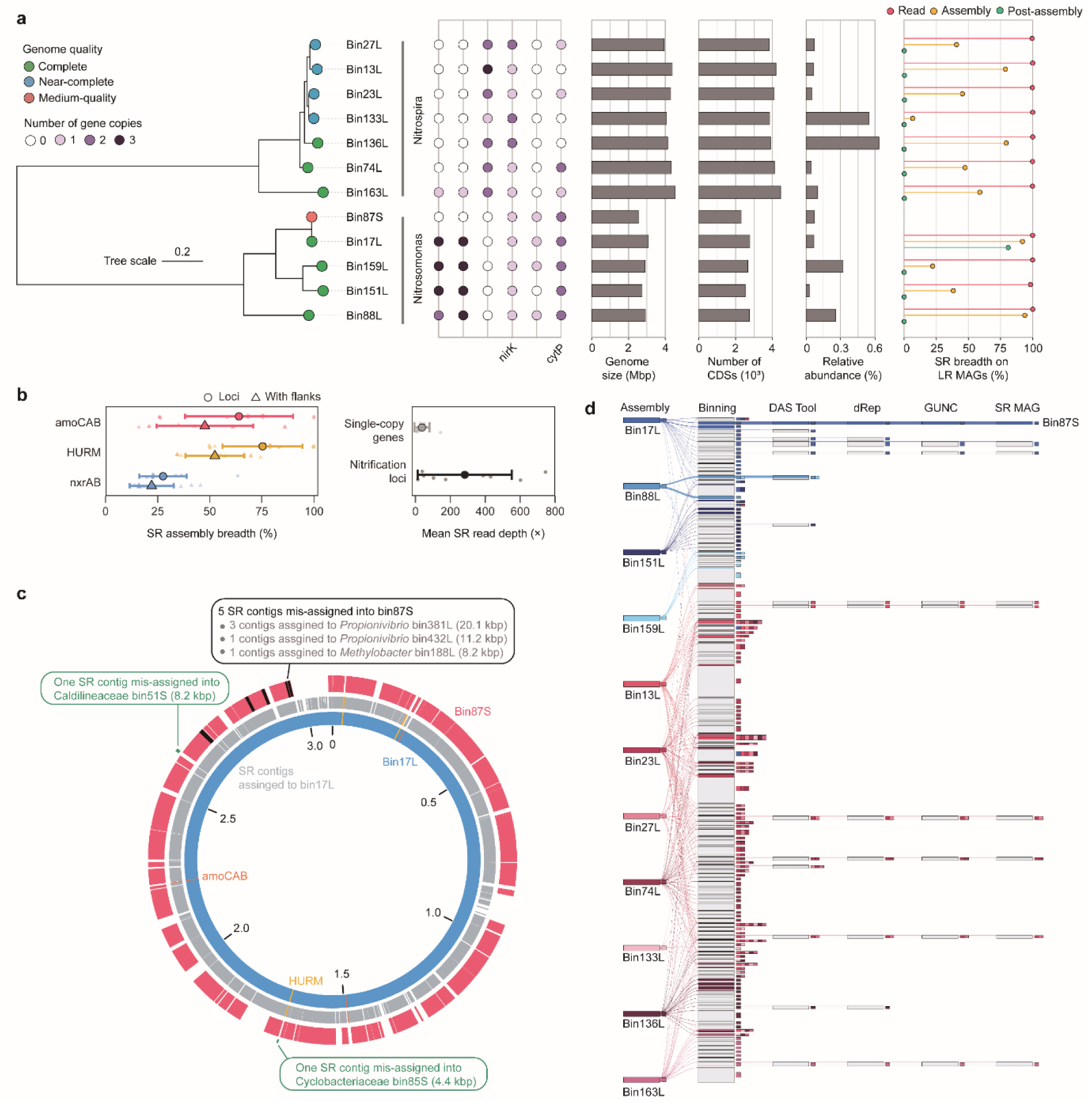
Comparative characterization of SR and LR nitrifier MAGs and mechanistic analysis of incomplete SR genome recovery. **a,** Phylogenomic tree of nitrifier MAGs recovered using SR and LR metagenomics, accompanied by annotations of genome statistics and nitrification gene repertoires. **b,** SR assembly breadth of nitrification loci in LR MAGs and comparison of SR read coverage between nitrification loci and GTDB-defined single-copy genes. Point-range plots show means±s.d., and jittered points indicate individual loci. **c,** Comparison of bin17L, SR contigs assigned to bin17L (grey), and bin87S (red). Radial lines indicate start codon locations of *amoCAB* (orange) and HURM (yellow) loci. **d,** Stage-wise tracking of SR contigs assigned to nitrifier LR MAGs, from initial assembly to final MAG recovery. Node thicknesses are proportional to total contig size. Blue and red hues represent *Nitrosomonas* and *Nitrospira*, respectively. Grey blocks denote individual SR bins, and the adjacent coloured segments indicate co-binning of contigs assigned to different nitrifier LR MAGs. Flows terminating at intermediate stages indicate contigs excluded during genome reconstruction. Of the seven SR MAG candidates, only bin87S, conspecific with *Nitrosomonas* bin17L, was retained through the complete metagenomic workflow.

**Extended Data Fig. 4.**
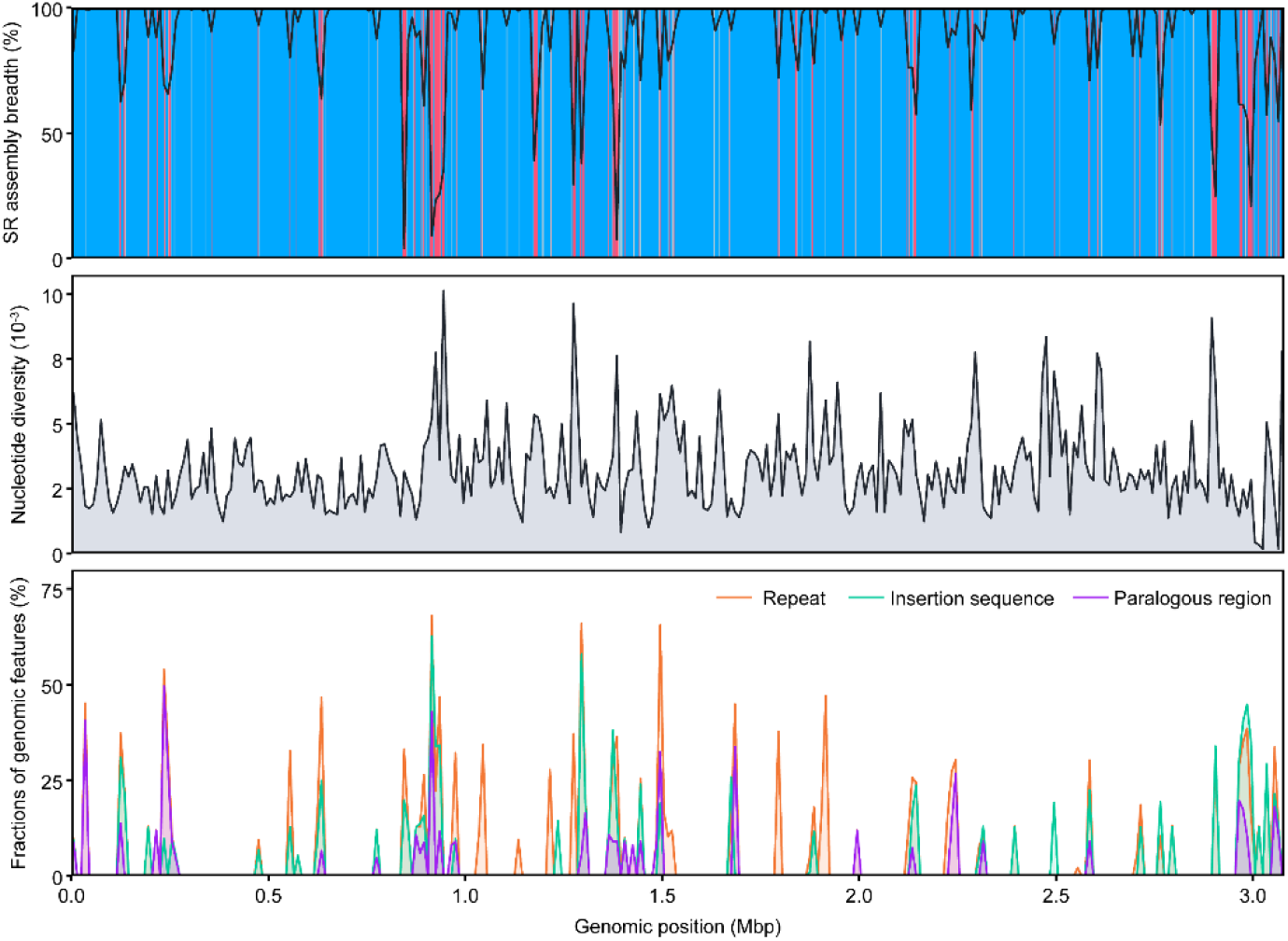
Genome-wide variation in SR assembly recovery (top), nucleotide diversity (middle), and frequencies of genomic feature (repeats, insertion sequences, paralogous regions, and provirus; bottom) along the complete genome of *Nitrosomonas* bin17L. Blue and red shading indicate local regions of near-complete (≥95%) and poor (<50%) SR assembly recovery, respectively. Recovery classes were defined in 1-kbp windows, whereas line profiles represent values smoothed over 10-kb windows.

In comparison, the SR workflow retrieved only a medium-quality *Nitrosomonas* MAG (bin87S) that was conspecific with bin17L (ANI=99.9%; AF= 80.4% for bin17L-to-bin87S and 97.4% for bin87S-to-bin17L). This MAG, consisting of 114 contigs (N50, 41,123 bp), exhibited a substantially reduced genome size (2.5 Mbp) and fewer protein-coding sequences (2,300 CDS), as compared to its LR counterpart (Fig. 4a). Importantly, *amoCAB* operons and the HURM module, whose presence is required to confirm the genomic potential for aerobic NH ^+^ oxidation, were absent from this SR genome. This omission of signature functional genes is a well-recognized limitation of analyzing nitrifying microbiomes with genome-resolved metagenomics^68,69^. Among 294 short-read AOB MAGs (213 *Nitrosomonas*, 78 *Nitrosospira*, and 3 *Ca. Nitrosoglobus*; average contig count, 269.0; mean N50, 85.6 kbp) publicly available as of 2026/03/26, only 19 genomes (17 *Nitrosomonas* and 2 *Nitrosospira*) encoded both at least one complete *amoCAB* operon and a complete HURM module (Supplementary Fig. 5; Supplementary Data 8).

Multiple lines of evidence support microdiversity as the main cause of the poor representation of nitrifier genomes in AS microbiome by SR metagenomics (Extended Data Fig. 4), although genome-level correlations were not statistically significant (Supplementary Note 10). Assembly breadth of SR contigs on LR MAGs were strongly biased towards bin17L (92.0%) and bin88L (93.9%), compared with 46.4±23.9% for the other nine nitrifier genomes (Fig. 4a; Supplementary Note 10; Supplementary Data 9). Nitrification marker loci in LR MAGs showed incomplete and fragmented recovery by SR assembly, particularly *nxrAB* loci in *Nitrospira* MAGs, which had an assembly breadth of 27.6±11.3%. Only five (three bin17L HURM and two bin88L *amoCAB* copies) were recovered with >90% coverage, and none together with their 1-kbp flanking regions, indicating that SR assembly constrained locus-specific recovery of these key functional elements in nitrifier genomes (Fig. 4b; Supplementary Data 10). The elevated read depth of the nitrification-related genes, relative to SCGs (282.5±269.2× versus 37.5±43.1×; *P*<0.05), supported homologous-copy collapse, also suggesting extensive microdiversity among the organisms harboring these genes (Fig. 4b).

Tracing the fate of the SR contigs assigned to the LR MAGs through the binning process identified cross-genome mixing as an additional cause of their poor representation in SR MAGs (Supplementary Note 11). The only qualified SR MAG bin87S contained 104 contigs assigned to its LR counterpart bin17L, covering 80.2% of the complete bin17L genome and representing 87.2% of all SR contigs assigned to bin17L. Among the SR contigs excluded from bin87S were a 3.5-kbp contig that aligned to part of one of the three *amoCAB* operons in bin17L and a 4.4-kbp contig carrying the shared, near-complete HURM sequence, which was misbinned to Cyclobacteriaceae-affiliated bin85S (conspecific to HURM-lacking bin103L; Fig. 4c). This binning error causes an additional interpretive problem by falsely attributing genomic potential for NH_2_OH oxidation to the Cyclobacteriaceae organism.

The failure trajectories of the other nitrifier-affiliated SR contigs during binning clearly demonstrate that neither *SemiBin*-nor *MetaBAT*-based binning is effective at recovering nitrifier genomes from the SR assemblies (Supplementary Note 11). A *Nitrosomonas*-affiliated candidate SR bin, which covered 88.5% of bin88L, was nevertheless excluded during dereplication owing to 55.3% contamination (Fig. 4d). Two candidate *Nitrospira* SR bins showed extensive cross-genome mixing, with contigs assigned to seven and six LR *Nitrospira* genomes, respectively, without discernible patterns. Nine contigs (N50=4.5 kbp) assigned to five LR *Nitrospira* genomes persisted in six unrelated SR MAGs.

Genome-wide depth heterogeneity, homologous-copy collapse at nitrification loci, and cross-genome mixing during binning, all consistent with the difficulty in resolving nitrifier population microdiversity, contributed to incomplete nitrifier recovery at different stages of the SR workflow^70–72^. As these limitations have little relevance to the sequencing depth, they are unlikely to be overcome merely by increasing short-read sequencing depth; our findings underscore the value of HiFi long-read metagenomics for nitrification-focused genome-resolved analyses.

### Genome completeness governs metagenomic identification and functional inference of phosphorus-accumulating organisms

Processing of HiFi LR metagenome sequences recovered 29 MAGs (7 complete, 10 near-complete and 12 medium-quality) affiliated with previously identified genera of polyphosphate-accumulating organisms (PAOs), including 6 *Azonexus*^73^ (2.0±0.4% cumulative relative abundance), 12 *Ca.* Accumulibacter^73,74^ (1.6±0.2%), 3 *Methylophosphatis*^23^ (0.8±0.1%) and 8 *Phosphoribacter*^73,75^ (1.8±0.7%) genomes (Fig. 5). Twelve (near-)complete genomes, including 7 *Accumulibacter*, 2 *Azonexus*, and 3 *Methylophosphatis*, encoded the full gene repertoires essential for polyphosphate turnover and the synthesis/depolymerization of canonical storage compounds (e.g. polyhydroxyalkanoates and glycogen; Supplementary Data 7). The five (near-)complete *Phosphoribacter* genomes represented a notable exception, uniformly lacking the genetic potential for polyhydroxyalkanoate biosynthesis despite their high genome completeness. In contrast, the medium-quality MAGs affiliated with the same putative PAO taxa from both LR and SR workflows, except for the *Accumulibacter* MAG bin248L, lacked at least one of the key markers. These genomic observations provide evidence, albeit indirect, that the previously reported gaps in the functional gene inventories for these PAO taxa reflect incomplete genome recovery rather than genuine metabolic variation^23,76^. The exceptional case of *Phosphoribacter* MAGs was consistent with previous genomic and *in situ* observations showing that *Phosphoribacter* can accumulate and cycle polyphosphate, suggesting a non-canonical PAO metabolism in this lineage^75^.

**Fig. 5:**
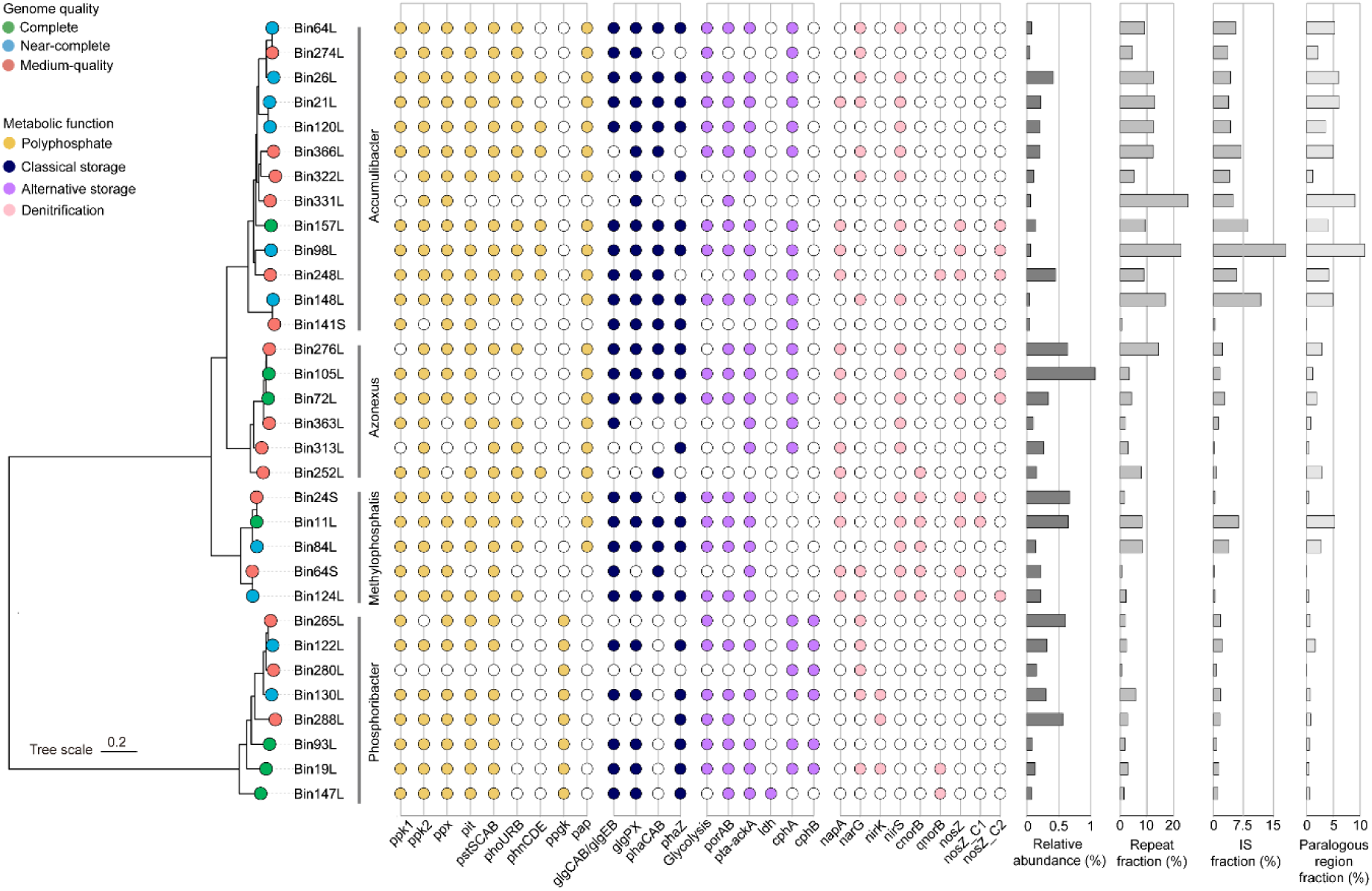
Phylogenomic tree of canonical PAOs recovered by LR and SR metagenomics. Tree tip colors indicate genome quality, and the adjacent strip indicates genus-level affiliation. The dot matrix shows the presence (filled circles) or absence (open circles) of genes encoding metabolic functions associated with enhanced biological phosphorus removal. The adjacent bar plots show mean relative abundance, and genomic fractions of repeats, insertion sequences (IS) and paralogs.

Close examination of the architectures of the seven (near-)complete single-contig *Accumulibacter* MAGs suggested insights as to why genome closure remains particularly challenging for this uncultivated lineage, even with HiFi LR metagenomics^23,77^. These genomes contained abundant repetitive sequences (13.7±4.8% of the genome length), insertion sequences (8.0±5.3% of the genome length) and paralogous genes (5.8±2.4% of all coding sequences), consistent with previous reports of *Accumulibacter* genome reconstruction from metagenomes (Supplementary Note 12)^77–79^. Such repetitive and multicopy sequences are both well-established constraints on chromosome closure in LR assemblies^6^; however, this analysis places particular emphasis on the abundance of repetitive regions. Repetitive regions occurred within 1 kb of 8 termini in five of six near-complete (unclosed) genomes, demonstrating that the extensive repeats barred contiguous assembly. The only complete genome bin157L, closure of which was supported by 12 HiFi reads directly spanning the end-to-start junction, was free of repetitive regions within 1 kb of either terminus (Supplementary Fig. 6; Supplementary Data 11). This contrast suggests that the extensive repetitive sequences represent a key mechanistic difficulty in obtaining complete genomes of this elusive taxon.

## Conclusion

The performance of biological nitrogen removal and enhanced biological phosphorus removal in modern-day WWTPs depends largely on the establishment and maintenance of balanced and resilient microbiomes with members contributing to nitrification, denitrification, and phosphorus accumulation. Through an exhaustive comparison between genome-resolved metagenomic analyses using conventional Illumina short-read shotgun sequencing and HiFi long-read sequencing, with particular emphasis on key functional guilds, we show that current genomic interpretations of the key

AS processes can be substantially distorted by incomplete and biased genome reconstruction. The advance enabled by LR metagenomics lies not merely in the number or diversity of closed and near-complete genomes recovered. Rather, we show that SR metagenomics substantially distorts estimates of the abundance and functional potential of ammonia oxidizers, nitrite oxidizers, and comammox populations that underpin the oxic segment of nitrogen removal in WWTPs, whereas HiFi LR metagenomics largely resolves these biases. While HiFi LR metagenomics provided high-confidence genomic evidence for the predominance of modular over complete denitrifiers, the comparisons show that conclusions regarding overall denitrification performance and N_2_O emissions based on SR denitrification gene inventories are prone to substantial biases. In addition, the expanded set of

(near-)complete genomes for uncultivated putative phosphorus-accumulating organisms provided crucial genomic support for previously proposed physiological mechanisms. Most importantly, using LR MAGs as references for exhaustive analyses of SR assembly and binning, we present mechanistic evidence that the deficiencies of genome-resolved metagenomics arise from intrinsic genomic features, rather than insufficient sequencing coverage, underscoring the need to incorporate HiFi LR metagenomics into AS microbiome analyses.

## Data availability

The raw DNA reads and 182 (near-)complete MAGs have been deposited in the NCBI Sequence Read Archive and GenBank, respectively, under BioProject accession number PRJNA1552241. All 447 long-read and 141 short-read MAGs were deposited in Zenodo (https://doi.org/10.5281/zenodo.23755631).

## Code availability

The workflow for assessing MAG assembly integrity is publicly available on GitHub at https://github.com/songmj86/MAGverify.

## Acknowledgement

This work was supported by the National Research Foundation of Korea (NRF) grant funded by the Korea government (MSIT) (No. RS-2024-00337570; RS-2026-25525273).

## Contributions

M.J.S. conceptualized the study and performed the data analyses. S.Y. supervised the study and guided its direction. T.Y. and J.Y.L. conducted sample preparation for shotgun metagenomic sequencing. T.Y., S.Y., and J.Y.L. supported the data analysis. M.J.S. wrote the initial manuscript draft, which was then extensively rewritten and revised by S.Y. A.J.P. critically reviewed and commented the manuscript. All authors reviewed and approved the final manuscript for submission.

