## Supplementary Information for "HiFi long-read metagenomics reshapes genome-resolved interpretations of activated sludge microbiome"

Corresponding author:

**Supplementary Methods.**

**HiFi read mapping for assembly validation.** HiFi-read validation was based on a modified version of the read-to-assembly framework of Trigodet et al[1]. Pooled HiFi reads were separately mapped to each assembly using *Minimap2* v2.31 (-ax map-hifi -p 1 --secondary-seq), and alignments were sorted and indexed using *SAMtools* v1.21. All mapped alignments were used for read-clipping, phantom-sequence and assembly-variant analyses, whereas only primary and supplementary alignments were used for all read-spanning analyses (circularity, false closure, false duplication and the spanning-support analyses below). Analyses were conducted at the contig level and summarized at the MAG level unless otherwise specified.

To assess end-to-start circular topology, each single-contig MAG was rotated at its midpoint to relocate the original end-to-start boundary internally, following the principle underlying *Circlator*[2]. HiFi reads were remapped to the LR assembly containing the rotated MAG contig using the *Minimap2* map-hifi preset, and junction support was assessed from primary or supplementary alignments. An alignment was accepted as spanning support only when it crossed the junction without deletions or skipped segments and extended ≥2 kbp on both sides. MAGs were classified as having HiFi-supported circular topology when at least two distinct HiFi reads spanned the end-to-start junction. MAGs were considered to have high-confidence circularity only when they were both assembler-circular (reported as circular by the assembler) and HiFi-supported.

To detect read-clipping sites and false closures, candidate sites were identified using *anvi-script-find-misassemblies* in *anvi'o* v9, with a clipping ratio >0.8 and the minimum distance to contig ends set to 0 bp[3]. A site was classified as HiFi-supported clipping when at least two distinct HiFi reads were clipped at the site and no HiFi read spanned the position with aligned bases on both sides. Sites within 100 bp of either contig end were classified as terminal, and the remainder as internal. Sites with ≥10× coverage and a clipping ratio of 1.0 were additionally designated high-confidence clipping sites, following Trigodet et al[1]. Single-contig MAGs were classified as premature-circularization candidates when a clipping site on an assembler-circular contig showed false closure, defined as cross-contig connections from at least two distinct clipped HiFi reads on each side. As an additional screen, circular contigs <500 kbp with at least three distinct ribosomal-protein families, identified using the anvi'o Bacteria_71 and Archaea_76 HMM collections, were flagged as suggestive of premature circularization[1].

To identify phantom sequences (assembled sequences lacking HiFi-read support), HiFi read depth was calculated using *SAMtools* depth (-aa -g SECONDARY). Consecutive zero-depth positions were merged, and intervals >500 bp were searched against HiFi reads using BLASTN (-dust no -max_target_seqs 1)[4]. Regions containing no ambiguous bases and lacking BLASTN hits were classified as phantom-sequence candidates. For validation, *Meryl* v1.4.1 was used to quantify the fraction of distinct 21-mers within these regions absent from HiFi reads[5].

To quantify repetitive sequences, non-diagonal self-alignments within contigs and alignments between contigs of the same genome were identified using *BLASTN* (-task blastn), retaining matches >200 bp with ≥80% nucleotide identity. Repeat fractions were calculated from merged, non-overlapping repeat intervals, using intra-contig repeats for single-contig MAGs and both intra- and inter-contig repeats for multi-contig MAGs. Contigs and genomes with repeat fractions ≥70% were classified as excessive-repeat[1].

To assess taxonomic chimeras, CDSs predicted from LR contigs using *Prodigal* (meta mode) were classified using the *taxonomy* module of *MMseqs2* v17.b804f (--lca-mode 3) against GTDB R232[6]. Each gene boundary was assessed at domain-to-order levels using up to five classified genes selected from the ten nearest genes on either side, with ≥80% agreement required to assign a dominant taxon. Discordant dominant taxa indicated a candidate transition. Only transitions overlapping a HiFi-supported clipping site were classified as chimeras, with those meeting the high-confidence clipping criterion flagged separately[1].

To assess false duplication, LR assemblies were self-aligned using *Minimap2* (-x asm5 -DP), retaining candidate pairs ≥1,000 bp with ≥95% sequence identity. Same-orientation alignments fully nested within another at both loci or showing ≥90% reciprocal overlap were merged. False duplication was identified when both boundaries of one copy were spanned by at least two distinct HiFi reads extending ≥500 bp into each flank, whereas neither boundary of the other copy was spanned despite equivalent flank support.

To assess minority and unsupported assembly variants, nucleotide and indel support was profiled using *anvi-profile* in *anvi'o* v9[3]. An SNV was classified as a minority assembly variant when assembly nucleotide retained HiFi-read support but an alternative nucleotide was supported by >50% of aligned reads, and as unsupported when the assembly nucleotide lacked HiFi-read support. Indels were classified as unsupported when indel counts equalled coverage and as minority when counts were >50% but below coverage. Variant counts were expressed per Mbp of the contig length.

**Reference-guided assessment of short-read genome recovery.** Paired SR reads were competitively mapped to the 447 LR MAGs using *Minimap2* (-ax sr -p 1). Coverage breadth at ≥1× and mean read depth were calculated from primary alignments using *BEDTools genomecov*, including zero-coverage reference positions. Depth heterogeneity was defined as the coefficient of variation of mean depths across non-overlapping 1-kbp windows. Primary SR read alignments were profiled using *inStrain* v1.10.0 (--min_mapq 2 --min_genome_coverage 1) to quantify nucleotide diversity and SNV density[7]. For local analyses, non-overlapping 1-kbp windows were defined along each LR contig, and SR read support, microdiversity and fractions overlapping repeats, insertion sequences, paralogous regions, and proviruses were quantified.

SR contigs were competitively aligned to the LR MAG catalogue using *Minimap2* (-x asm5 -c --cs=long -N 50). Alignments with ≥99% nucleotide identity and ≥2.5 kbp aligned length were retained, and overlapping alignments to the same LR MAG were merged. For each SR contig, the best-supported LR MAG was selected based on total aligned length, with length-weighted nucleotide identity used to resolve ties. Contigs with <65% alignment breadth or equally supported best matches to multiple LR MAGs were left unassigned.

To assess fragmentation, regions between consecutive SR contig alignments along each LR MAG not bridged by a single SR contig were classified as gapped when intervening reference sequence remained uncovered and as gapless when the alignments were directly adjacent. For gapped regions, the lengths of coding and non-coding sequences, repeat, insertion-sequence, provirus and RNA-feature sequence were measured. Candidate misassemblies in assigned SR contigs were identified using *QUAST* v5.3.0 (≥500 bp alignment length and ≥99% nucleotide identity)[8]. Fragmented regions and QUAST-reported junctions were considered supported when at least two distinct HiFi reads spanned them with ≥500 bp aligned on both sides.

To assess contig retention during post-assembly processing (Methods), the number, total length and assembly breadth of SR contigs assigned to each LR MAG were quantified at each stage. Contigs were classified as unbinned, lost during processing or incorporated into a final SR MAG. The proportion of sequence in each final SR MAG assigned to non-paired LR MAGs was also quantified.

**Genome analysis.** CRISPR arrays and spacers were identified in 447 LR and 141 SR MAGs using *CRISPRCasTyper* v1.8.0 (--prodigal single)[9]. Microbial lifestyles were predicted using *SymClatron* v0.10.10 from *Prodigal*-predicted protein sequences, retaining only classifications passing the confidence threshold[10]. Microbial growth potential was estimated from genomic codon usage patterns using *gRodon2* v2.5.2[11]. Genes annotated as 30S or 50S ribosomal proteins were designated as highly expressed genes, excluding annotations associated with methylation or hydroxylation. Predicted minimum doubling times were calculated using the *predictGrowth* function in full and partial mode for (near-)complete and remaining MAGs, respectively.

**Orthogroup-level comparisons of conspecific genomes.** For each genome pair, concordance and discordance were calculated as the proportions of orthogroups with identical or differing copy numbers between MAGs, respectively. Discordant orthogroups were classified by copy-number excess or exclusive presence in either MAG. For orthogroups with higher LR copy numbers, paralog pairs were assessed with *MMseqs2 easy-search*[6], with ≥70% amino acid identity and ≥70% alignment coverage for both proteins indicating close relatedness. For orthogroups with higher SR copy numbers or exclusive to SR MAGs, SR proteins were aligned to the paired LR genome using *miniprot* v0.18 (-S --outs=0.95 --outc=0.20)[12]. Alignments with ≥80% alignment fraction and ≥90% identity were used to identify potential gene fragmentation underlying SR copy excess.

**Targeted analyses of nitrifiers and Ca. *Accumulibacter*.** Eleven (near-)complete nitrifier LR MAGs (four *Nitrosomonas*, six canonical *Nitrospira* and one comammox *Nitrospira*) were examined at the genome level and locus level across 39 nitrification loci (12 *amoCAB*, 13 HURM and 14 *nxrAB*). SR read breadth and mean depth were calculated for these loci and their 1-kbp flanks. LR locus and flanking sequences were aligned to the SR assembly using *Minimap2* (-x asm5 -c --cs=long), with merged alignments used to assess coverage, nucleotide identity and fragmentation. Locus depth was compared with that of bacterial SCGs (single-copy GTDB-Tk bac120 markers) in two coordinate systems. In SR-assembly coordinates, SR contig regions onto which ≥2 LR copies of the same locus aligned (collapsed regions) were compared with SCGs on the SR contigs assigned to the same LR MAG, using SR reads mapped to the SR assembly; in LR MAG coordinates, the same loci were compared with SCGs on each LR MAG, using SR and HiFi reads competitively mapped to the 447 LR MAGs. Depth was calculated from primary alignments as mean per-base depth over each locus or gene, averaged per MAG. For seven *Ca.* Accumulibacter LR MAGs, repeats, insertion sequences and paralogs with at least 1-kbp flanks were classified as spanned when at least two distinct HiFi reads covered both flanks and ≥90% of the feature.

**Statistical analysis.** Wilcoxon rank-sum tests compared SR read support, assembly recovery and sequence retention between nitrifier and non-nitrifier MAGs, whereas Kruskal-Wallis tests assessed these metrics across *Nitrosomonas*, canonical *Nitrospira* and non-nitrifier MAGs.

### Supplementary Notes

**Supplementary Note 1. Long-read sequencing improved assembly contiguity and gene integrity.**

LR sequencing generated substantially longer reads than SR sequencing (16.2 kbp versus 151 bp), with comparable Q30 fractions (93.4±0.4% versus 93.3±0.1%; Supplementary Table 2). The three LR assemblies yielded 10.7±2.0 Gbp, ~55% more assembled sequence than the SR assembly (6.9 Gbp). LR assemblies were markedly less fragmented, with fewer contigs (206,304±114,310 versus 12,619,252), longer mean (59.3±21.1 kbp versus 549.4 bp) and maximum (10.2±1.5 Mbp versus 0.6 Mbp) contig lengths, and higher N50 (79.1±24.9 kbp versus 771 bp). Circular contigs occurred only in LR assemblies (0.94±0.06% of contigs; 1,108 to 3,218 per assembler). Although the SR assembly yielded more gene predictions than LR assemblies (16.0 versus 10.5±2.2 million), only 13.2% contained both start and stop codons versus 97.0±1.3% in LR assemblies. Consequently, LR assemblies yielded nearly fivefold more complete genes (10.1±2.1 versus 2.1 million).

**Supplementary Note 2. HiFi reads support the assembly integrity of long-read MAGs. Assembly integrity was evaluated in 447 LR MAGs, with circularity assessed in 350 single-contig MAGs (Supplementary Data 2). Of these, 107 showed HiFi-supported circularity based on at least two distinct junction-spanning HiFi reads, whereas 135 lacked such support, and 108 were not evaluable. Of the 103 assembler-circular MAGs, 102 (99.0%) met the criterion and were classified as high-confidence circular genomes. Five additional MAGs showed HiFi-supported circularity without assembler-reported circularity.**

**High-confidence clipping was rare.** HiFi-supported clipping was detected at 114 sites across 65 MAGs (47 single-contig and 18 multi-contig MAGs), including 111 internal and three terminal sites. Of these, 100 occurred in *metaMDBG* MAGs, 13 in *metaFlye* MAGs and one in a *hifiasm-meta* MAG, consistent with the assembler-dependent pattern reported by Trigodet et al[1]. Most MAGs (55 of 65; 84.6%) showed one or two sites, with a maximum of seven in bin132L. Only nine sites in seven MAGs met the high-confidence clipping criterion (100% clipping at ≥10× HiFi coverage), comprising eight sites in six *metaMDBG* MAGs (bin26L, bin41L, bin51L, bin262L, bin339L and bin359L) and one site in the *metaFlye* MAG bin335L. Twelve of the 103 assembler-circular MAGs contained 18 internal HiFi-supported clipping sites, none showing bilateral HiFi-supported cross-contig connections indicative of false closure. Bin320L met the ribosomal-protein proxy criterion for circular MAGs shorter than 500 kbp but showed no HiFi evidence of false closure[1]. No single-contig MAG was classified as a premature-circularization candidate, and seven assembler-circular MAGs classified as medium quality under MIMAG are discussed in Supplementary Note 3.

Neither excessive repeats nor false duplications were detected. Repeat regions were detected in all MAGs except bin320L (157.5±196.5 kbp per genome; 5.7±7.3% of genome length), with 65 MAGs (14.5%) showing genome-wide repeat fractions ≥10%. The highest repeat fraction was observed in bin443L (690.4 kbp; 43.7% of its 1.6-Mbp genome), while the largest total repeat length was detected in the *Accumulibacter*-affiliated bin98L (1.4 Mbp; 22.9% of its 6.0-Mbp genome). The 70% excessive-repeat threshold proposed by Trigodet et al. was not reached at either the contig or the genome level[1]. False-duplication screening identified 12,032 candidate pairs in 308 MAGs, comprising 11,710 intra-contig and 322 inter-contig pairs. Of these, 11,663 candidate pairs (96.9%) showed HiFi-read support at their boundary and none met the false-duplication criterion.

Phantom sequences and taxonomic chimeras were rare. Eleven regions lacking HiFi-read support were detected in 10 *metaMDBG* MAGs (0.02% to 0.14% of genome length). These comprised one region (505 to 2,574 bp) in each of nine MAGs and two in bin37L (3,914 bp in total). Meryl analysis showed that 13,105 of 13,488 distinct 21-mers (97.2%) in these regions were absent from HiFi reads. Six MAGs (bin18L, bin36L, bin37L, bin70L, bin73L and bin132L) were reclassified as high quality after phantom-sequence screening, whereas four (bin99L, bin231L, bin290L and bin444L) remained medium quality owing to other genome-quality criteria. For taxonomic chimera assessment, gene-level taxonomic transitions were detected in 223 MAGs (49.9%), but only two MAGs (bin197L and bin442L) showed transitions coinciding with HiFi-supported clipping sites and met the taxonomic chimera criterion, although neither met the high-confidence clipping criterion of Trigodet et al[1]. These involved transition from Gemmatimonadota to Actinomycetota in bin197L and from Desulfobacterota to Bdellovibrionota in bin442L.

Minority variants accounted for most assembly variants. Among 702,005 variants (533.7 per Mbp), 690,400 (98.3%) across 418 MAGs (93.5%) were classified as minority variants, with HiFi-supported assembly alleles less abundant than alternative alleles. The remaining 11,605 variants (1.7%) across 275 MAGs (61.5%) were unsupported variants, with no HiFi read supporting the assembly allele. Minority variants were interpreted as read-to-assembly allele disagreements rather than definitive assembly errors, as they may reflect within-population sequence variation[7].

**Supplementary Note 3. Circularity does not guarantee genome completeness.** Seven assembler-circular MAGs remained medium quality under MIMAG despite HiFi-supported circularity and no evidence of false closure (Supplementary Note 2). Of four *Promineifilum* MAGs, three (bin33L, bin58L and bin233L) lacked complete rRNA operons, whereas bin227L and bin233L showed estimated completeness below 90%. The multipartite reference genome of *Candidatus Promineifilum breve* raised the possibility that the three rRNA-lacking MAGs represented secondary chromosomes[13]. However, their sizes (4.1±0.1 Mbp) were closer to chromosome I (4.0 Mbp) than to rRNA-lacking chromosome II (1.0 Mbp), with limited similarity to chromosome I (ANI, 78.5 to 82.0%; AF, 20.5 to 42.3%). Read tracing identified 107 *Promineifilum*-affiliated 16S rRNA reads (98.7±0.8% nucleotide identity; 1,453±29 bp alignment length) and confirmed their recovery in assembled contigs. However, rRNA-encoding contigs were found in only two bins (3.4 Mbp each), neither of which was retained during dereplication in favor of longer representatives. Only bin33L was assigned to a GTDB species whose representative encoded a complete rRNA operon, but no bin33L sequence aligned to that operon. Together, genome comparisons did not support a secondary-chromosome origin, while read tracing confirmed that *Promineifilum* 16S rRNA sequences were assembled but not incorporated into three MAGs.

Among the remaining three MAGs, Patescibacteriota bin175L lacked a complete 16S rRNA gene, whereas Obscuribacteraceae bin141L marginally exceeded the 5% contamination threshold (5.03%). *Azoamicus* bin320L comprised a 246.8-kbp contig (20.6% GC content; 63 SCGs) and was predicted to be an obligate intracellular symbiont, similar to the reduced-genome ciliate endosymbiont *Ca. Azoamicus ciliaticola*[14]. Although completeness may be underestimated in reduced *Azoamicus* lineages (90.1% for the complete *Ca. A. ciliaticola* genome), the lower estimate for bin320L (80.8%) suggests incomplete genome recovery.

**Supplementary Note 4. Long-read metagenomics resolves genomes with unusual sizes and architectures.** LR metagenomics recovered high-quality genomes of understudied but potentially ecologically significant community members by resolving genomes with atypical sizes and complex architectures. Members of *Patescibacteriota*, a phylum characterized by unusually small genomes, were recovered *en masse*, comprising 15 complete, 17 near-complete, 3 high-quality, and 45 medium-quality MAGs with a cumulative relative abundance of 1.8±0.4% (Supplementary Fig. 1; Supplementary Data 1)[15]. The complete and near-complete genomes were 1.0±0.2 Mbp in size, showcasing the hallmark features of genome streamlining, consistent with symbiotic lifestyles[10]. Their reductive evolution likely minimizes functional redundancy, as all (near-)complete MAGs encoded relatively few single-copy marker genes (89.6±6.8), with eleven containing no paralogous genes in the entire genomes. Unlinked rRNA architecture, previously reported only in unbinned long-read sequences for *Patescibacteriota*, was found in 15 (near-)complete MAGs[16].

At the opposite end of the spectrum, LR metagenomics reconstructed a near-complete Myxococcota MAG (bin87L) as a single 11.1-Mbp contig. Genomes of comparable size have been reported for cultivated myxobacteria, including *Chondromyces crocatus* and *Sorangium cellulosum*, but, to our knowledge, not as a single-contig MAG with independently supported contiguity[17-19]. Such high-confidence contiguity is particularly crucial for verification of large-sized genomes, as unresolved assembly or binning artifacts could otherwise inflate genome-size estimates. Genome size within *Myxococcota* can vary substantially, as illustrated by another near-complete MAG (bin115L), which shared 83.1% genome-wide ANI and 85.8% 16S rRNA gene identity with bin87L but possessed a markedly smaller genome of 2.2 Mbp. Together, the *Patescibacteriota* and *Myxococcota* examples demonstrate the importance of highly contiguous genome reconstruction for robust inference of extreme genome sizes from metagenomic data.

**Supplementary Note 5. Long-read metagenomics reveals genomic complexity obscured by short-read assemblies.** Genome reconstruction and feature recovery were compared across 56 conspecific genome pairs (ANI, 99.8±0.2%; ANI, AF, 81.5±8.3% of the LR genome and 95.3±5.2% of the SR genome aligned), pairing 35 complete and 21 near-complete LR MAGs with medium-quality SR MAGs under the MIMAG framework (Supplementary Fig. 3; Supplementary Data 4). SR MAGs were highly fragmented (234.2±119.7 contigs per genome), with mean N50 values >100-fold lower than those of LR MAGs (31.1±23.6 versus 4,152.3±1,121.9 kbp; Wilcoxon signed-rank test, $P<0.05$). LR MAGs also showed higher completeness (98.3±2.4% versus 90.2±6.9%) and lower contamination (0.8±0.9% versus 1.7±1.7%; both $P<0.05$), and 24 SR MAGs failed MIMAG high-quality threshold for completeness (>90%) or contamination (<5%). Complete rRNA operons were detected in all LR MAGs, whereas no SR MAG encoded a 16S rRNA gene. SR MAGs encoded 17.3±2.2 distinct tRNAs on average, with 27 below the MIMAG threshold of 18.

LR metagenomics improved the recovery of protein-coding genes and structurally complex genomic regions. LR MAGs encoded more CDSs (3,713±824 versus 3,328±769) and a higher fraction of complete CDSs (99.95±0.03% versus 90.0±5.3%; both $P<0.05$). Repetitive sequences (157.7±117.6 versus 34.6±55.8 kbp per genome), insertion sequences (34.8±42.0 versus 6.1±6.4 per genome) and paralogous genes (43.9±50.0 versus 4.6±15.5 per genome) were also more abundant in LR MAGs (all $P<0.05$). LR MAGs recovered more proviruses (36 in 18 LR MAGs versus 8 in 7 SR MAGs) and CRISPR arrays (67 in 28 LR MAGs versus 21 in 13 SR MAGs; both $P<0.05$). Screening 2,707 CRISPR spacers from LR MAGs and 170 from SR MAGs against 260,256 IMG/VR viral genomes identified more virus-matching spacers in LR MAGs (81 in 15 LR MAGs versus 7 in 3 SR MAGs), indicating a broader set of potential host-virus associations.

**Supplementary Note 6. Conspecific genome pairs differ in gene-family content.** Orthogroups (OGs) were compared across 56 conspecific genome pairs for gene-family content and copy-number differences, with multiple genes from the same genome assigned to an OG treated as paralogs (Supplementary Fig. 4; Supplementary Data 5). Of 174,393 OG comparisons (3,114.2±718.3 per genome pair), 161,210 showed concordant copy numbers (2,878.8±698.5 OGs; 92.4±3.9% of total OGs per pair), with only 15 of 56 genome pairs exceeding 95% concordance. Copy-number discordance occurred in 13,183 comparisons (235.4±131.0 OGs; 7.6±3.9% per pair), with the lowest copy-number concordance in the Microtrichaceae-affiliated bin24L-to-bin68S pair (81.7%). Of these, 7,571 were shared OGs with higher copy numbers in LR MAGs (135.2±83.9 OGs; 57.1±15.2% of discordant OGs per pair) and 1,895 were LR-only OGs (33.8±27.8 OGs; 15.0±8.5% per pair of discordant OGs per pair), together accounting for 71.8% of all discordant OG comparisons. Paralog similarity within shared OGs was assessed to determine whether higher LR copy numbers primarily involved closely related paralogs potentially collapsed during SR assembly[20]. Only 21.4% of 16,293 pairwise alignments showed ≥70% amino-acid identity and ≥70% bidirectional AF, indicating that LR copy excess was not restricted to closely related paralogs. In contrast, 3,459 shared OGs showed higher copy numbers in SR MAGs (61.8±59.1 OGs; 26.2±14.6% per pair of discordant OGs per pair), and 258 were SR-only OGs (4.6±8.1 OGs; 1.7±2.2% per pair of discordant OGs per pair). To assess whether fragmented SR gene predictions inflated SR copy numbers, SR genes were mapped to paired LR MAGs using *miniprot*[12]. Following homologous-locus reassessment, 3,354 of 3,459 OGs (97.0%) retained higher SR locus counts, whereas LR homologs were identified for only 34 of 258 SR-only OGs (13.2%). Most SR-biased differences were therefore not explained by fragmented gene prediction and may involve strain-level gene-content variation or differential assembly and binning of closely related strains[21].

**Supplementary Note 7. Short-read assembly is the primary bottleneck in genome recovery.**

Quality-filtered SR reads covered nearly the entire length of each LR MAG (99.3% to 100% at ≥1×; 99.8±0.2%), with mean depths ranging from 6.8× to 145.8× (24.7±25.6×; Supplementary Data 6). In contrast, at the assembly level, SR contigs assigned to each LR MAG covered 50.2% to 98.4% of LR MAG length (84.0±10.6%), with assembly breadth subsequently decreasing after initial binning (81.8±11.0%), bin consolidation (76.8±12.0%) and in the paired SR MAGs (76.1±12.1%; Supplementary Data 6). Assigned SR contigs and paired SR MAGs were similar in total length (3.51±1.00 versus 3.55±0.98 Mbp; Wilcoxon signed-rank test, $P>0.05$) and contig number (250.5±128.7 versus 234.2±119.7), supporting the comparability of the two genome sets. Assembly-stage losses (16.0±10.6% of LR MAG length) exceeded post-assembly losses (7.9±4.3%) in 45 of 56 pairs.

HiFi reads revealed extensive fragmentation with limited misassembly. Across the 56 genome pairs, 10,364 fragmented regions (10,291 gapped and 73 gapless) were identified between consecutive SR contig alignments along LR MAGs (Supplementary Data 6). Protein-coding and non-coding sequences accounted for 84.6±6.6% and 15.4±6.6% of the total gapped-region length, respectively. Of the 10,291 gapped regions, 9,865 (95.9%) were spanned by at least two distinct HiFi reads extending ≥500 bp into both flanks. QUAST identified 51 candidate misassembly junctions (24 extensive and 27 local) across 34 conspecific genome pairs. Contigs carrying these candidates totaled 2.1 Mbp (61.8±58.4 kbp per genome; 2.2±3.0% of assigned SR contig sequence). Of the 51 candidate junctions, 43 (all 27 local and 16 extensive) were spanned by at least two distinct HiFi reads extending ≥500 bp into both flanks. HiFi support for most QUAST candidate junctions suggests that these reference-relative discrepancies may reflect genuine genomic variation rather than SR assembly errors[8].

During post-assembly processing, initial binning and bin consolidation resulted in losses of 2.2±1.4% and 5.0±3.7% of LR MAG length, respectively, with the largest decrease occurring during bin consolidation in 43 of 56 pairs. Most SR contig sequence assigned to LR MAGs was retained in paired conspecific SR MAGs (90.3±5.3%). Sequences assigned to non-paired LR MAGs accounted for only 1.6±2.8% of final SR MAG length (exceeding 5% in five of 56 MAGs). Consistently, greater decreases in assembly breadth during post-assembly processing were associated with lower SR assembly contiguity (partial Spearman's $\rho$ = -0.49) and higher SR contig numbers ($\rho$ = 0.37, both $P$<0.05). SR contigs not retained in paired MAGs also showed lower N50 values than retained contigs (7.7±4.1 versus 32.0±23.5 kbp per MAG; Wilcoxon signed-rank test, $P$<0.05).

**Supplementary Note 8. Microdiversity is linked to incomplete short-read assembly.** To identify factors associated with incomplete SR assembly, we examined read depth, depth variation, microdiversity and repetitive and mobile genomic features (insertion sequences and proviruses) in relation to assembly breadth and contiguity across 56 conspecific genome pairs (Supplementary Data 6). Microdiversity showed the strongest and most consistent associations, with nucleotide diversity and SNV density negatively correlated with assembly breadth (Spearman's $\rho$=-0.66 and -0.51, respectively) and contiguity ($\rho$=-0.55 and -0.38; all $P<0.05$). These associations persisted in partial Spearman analyses accounting for covariation among candidate factors, both for breadth ($\rho$=-0.70 and -0.64) and contiguity ($\rho$=-0.68 and -0.66; all $P<0.05$). Mean SR read depth was positively correlated with assembly contiguity in both Spearman ($\rho$=0.44) and partial Spearman analyses ($\rho$=0.38; both $P<0.05$), but not with assembly breadth ($\rho$=0.15 and 0.01, respectively; both $P>0.05$). SR read-depth heterogeneity was negatively correlated with assembly breadth in both Spearman ($\rho$=-0.33) and partial Spearman analyses ($\rho$=-0.32; both $P<0.05$), but not with contiguity ($\rho$=-0.24 and -0.09, respectively; both $P>0.05$). The fractions of repeats, insertion sequences and proviruses showed no significant associations with assembly breadth or contiguity in either analysis (all $P>0.05$). For local comparisons, 232,500 non-overlapping 1-kbp windows across 56 LR MAGs were classified by SR assembly breadth, with 31,743 showing poor (<50%) and 191,676 near-complete (≥95%) breadth. Read depth did not differ significantly between poorly and near-completely recovered windows (25.6±26.4× versus 24.8±25.6×; $P>0.05$).

**Supplementary Note 9. Comammox *Nitrospira* genomes share conserved metabolic features.**

Bin163L was compared with the complete genomes of *Nitrospira inopinata* and *Candidatus Nitrospira kreftii* to assess its genomic distinctiveness and conserved and lineage-specific metabolic features[22,23]. Bin163L was genomically distinct from *N. inopinata* (ANI, 85.9%) and *Ca. N. kreftii* (ANI, 83.7%), and its closest GTDB relative (*Nitrospira* sp. UW-LDO-01; ANI, 97.7%; AF, 55.2%) is represented by a fragmented draft genome (230 contigs; N50, 29.4 kbp). All three genomes encoded a single *amoCAB* operon, a HURM module, and the complete system I cytochrome c maturation machinery (*ccmABCDEFGHI*). The HURM and *ccm* genes occupied a conserved region in bin163L and *Ca. N. kreftii*, whereas the HURM module were separated from *amoCAB* in *N. inopinata*. All three genomes encoded copper-homeostasis genes (*copC* and *copD*) and bacterioferritin (*bfr*), suggesting shared potential for metal homeostasis and oxidative-stress protection[24]. Each also encoded several orphan *amoC*-like genes, which may contribute to stress recovery[25]. The *nxrAB* regions differed in number (one in *N. inopinata* versus two in bin163L and *Ca. N. kreftii*) and accessory-gene composition. Each genome encoded one *nirK* copy but lacked *norB*, *hmp* (flavohemoglobin) and *norVW* (flavorubredoxin). All three genomes encoded guanidine-utilization modules comprising a guanidine-I riboswitch (*ykkC*-*yxkD* class) upstream of an APC superfamily permease and guanidinase[26]. Complete urea-utilization machinery was also conserved, comprising the nickel-dependent urease (*ureABC*), the accessory genes (*ureDFG*) and the putative high-affinity urea transporter (*urtABCDE*), supporting the genomic potential for complete guanidine-to-$NO_3^-$ oxidation in all three genomes.

**Supplementary Note 10. Short-read assembly fragments nitrifier genomes and nitrification loci.**
Genome- and locus-level analyses focused on the 11 (near-)complete nitrifier LR MAGs, excluding five medium-quality *Nitrospira* LR MAGs (bin206L, bin255L, bin295L, bin329L and bin388L). Despite broad SR read support, the number of SR contigs assigned to them ranged from 40 to 506 (0.3 to 3.5 Mbp; Supplementary Data 9), with assembly breadth and contiguity (N50) comparable to the other 171 MAGs (54.9±28.5% versus 60.2±30.0%; 15.5±14.0 versus 16.2±17.6 kbp; Wilcoxon rank-sum tests, $P$>0.05). Canonical *Nitrospira* showed numerically lower values than *Nitrosomonas* (49.7±27.2% versus 61.6±36.8%; 10.3±6.6 versus 25.8±18.9 kbp; Kruskal-Wallis tests, $P$>0.05). Fragmentation patterns differed: many short gaps in bin163L, bin23L and bin74L (339 to 409; 5.7±1.1 kbp), fewer but longer gaps in bin133L and bin159L (38 to 57; 64.3±35.4 kbp), and only very short gaps in bin17L and bin88L (2.24±0.02 kbp). HiFi reads spanned 2,153 of 2,397 inter-contig gaps (89.8%) and the sole *QUAST*-reported discrepancy. Mean nucleotide diversity and SNV density were 39% to 70% higher in nitrifier MAGs than in other MAGs and twofold higher in canonical *Nitrospira* than in *Nitrosomonas* (all $P$>0.05), but, unlike conspecific genome pairs, neither microdiversity nor other factors were significantly associated with SR assembly breadth or contiguity (Spearman and partial Spearman, all $P$>0.05).

Nitrification-locus assembly breadth (54.7±28.3%; *amoCAB*, 64.0±25.9%; HURM, 75.3±19.2%; *nxrAB*, 27.6±11.3%) was comparable to genome-wide breadth (54.9±28.5%; $P$>0.05). One contig aligned to both bin88L *amoCAB* loci and another to all three bin17L HURM loci (identity and AF >99%), with similar multi-paralog alignments in 28 additional partially recovered loci. These collapsed regions showed higher read depth than SCGs (282.5±269.2× versus 37.5±43.1×; nine MAGs; $P$<0.05), but not in LR MAG coordinates (SR, 48.0±46.8× versus 48.6±58.7×; HiFi, 81.9±75.0× versus 90.5±99.5×; both $P$>0.05). Copy-specific context, including 1-kbp flanks, was incompletely recovered (86.0% for both bin88L *amoCAB* loci, 69.7% to 74.4% for the three bin17L HURM loci and 14.0% to 61.1% for the remaining 34 loci; Supplementary Data 10). HiFi reads supported 61 of 62 terminal fragmentation boundaries across 37 nitrification-related loci.

**Supplementary Note 11. Post-assembly processing limits nitrifier genome recovery.** The fraction of SR-assembly-covered LR MAG sequence retained in the final SR MAG catalogue was lower for nitrifier MAGs than for other MAGs (8.2±26.4% versus 33.7±43.2%; Wilcoxon rank-sum test, $P<0.05$), with the lowest mean retention in canonical *Nitrospira* (0.3±0.2% versus 22.0±44.0% in *Nitrosomonas*; Kruskal-Wallis test, $P>0.05$; Supplementary Data 9). Among the four AOB MAGs, 104 of 158 bin17L-assigned SR contigs (2.5 of 2.9 Mbp) entered the final bin87S through semibin2_64 (97.5% of bin87S and 87.2% of bin17L-assigned sequence). Consolidated semibin2_35 (4.5 Mbp) retained 79 bin88L-assigned contigs (2.6 Mbp), all of which were lost during dereplication. The two contigs carrying bin88L shared *amoCAB* and HURM loci remained unbinned. For bin151L and bin159L, SR assembly recovered 255 and 61 assigned contigs (38.2% and 22.2% breadth), of which 175 and 46, respectively, were retained by initial binners, mostly in heterogeneous candidate bins. Consolidation left one (4.4 kbp) and two (12.6 kbp) contigs, respectively, all of which were lost during dereplication.

For *Nitrospira*, MetaBAT_66 (1,386 contigs; 9.6 Mbp) and SemiBin2_172 (501; 5.6 Mbp) included 929 (6.8 Mbp) and 387 (4.3 Mbp) contigs from seven and six *Nitrospira* LR MAGs, respectively, but neither survived consolidation (53.4% and 27.0% contamination). Despite extensive SR assembly recovery for bin13L (358 contigs; 3.5 Mbp; 78.8% breadth) and bin136L (252; 3.4 Mbp; 79.4%), consolidation retained only two of 307 initially binned contigs (12.9 kbp of 3.3 Mbp) and one of 194 (7.0 kbp of 3.1 Mbp), respectively. The two bin13L-assigned contigs entered final bin90S (*Aestuariivirga*-affiliated; 8.5 kbp) and bin125S (Gammaproteobacteria-affiliated; 4.4 kbp), whereas the bin136L contig failed GUNC filtering. For the other five *Nitrospira* LR MAGs, SR assembly breadth ranged from 6.6% (bin133L) to 58.9% (comammox bin163L), with 23 to 325 assigned contigs initially binned per MAG but only three remaining per MAG after consolidation. Dereplication left seven contigs from four MAGs (three from bin23L, one each from bin27L and bin74L, and two from bin163L), all in non-conspecific final SR MAGs.

**Supplementary Note 12. Terminal repeats complicate genome closure in Ca. *Accumulibacter* MAGs.** Seven single-contig *Accumulibacter* LR MAGs were compared to assess previously reported constraints on complete genome recovery, including read depth, strain microdiversity, repetitive sequences and mobile genetic elements[27-29]. Repetitive regions occupied 9.1% to 22.9% of genome length, with individual region extending up to 84.2 kbp. The genomes also contained 105 to 609 insertion sequences, 152 to 587 paralogous genes (up to 54 copies per orthogroup) and one to four proviruses. Only bin157L met the high-confidence circularity criterion, although its sequencing depth, repetitive and mobile element content, paralogous gene fraction and strain microdiversity were comparable to those of the six near-complete genomes. Thus, these genome-wide characteristics did not clearly distinguish complete from near-complete recovery.

The clearest differences emerged at the contig termini. Among 12 contig termini of six near-complete genomes, repetitive regions occurred within 1 kb of eight termini across five genomes, whereas insertion sequences and paralogous genes each occurred within 1 kb of two termini (Supplementary Data 11). For bin64L, bin98L and bin120L, all six termini lay within 1 or 5 kb of a repetitive region. Repetitive regions and insertion sequences occurred within 1 kb of one end of bin21L, while large repetitive regions extended directly to both ends of bin26L. None of the six near-complete genomes yielded HiFi reads spanning the putative end-to-start junction with ≥2 kb alignment on both sides.

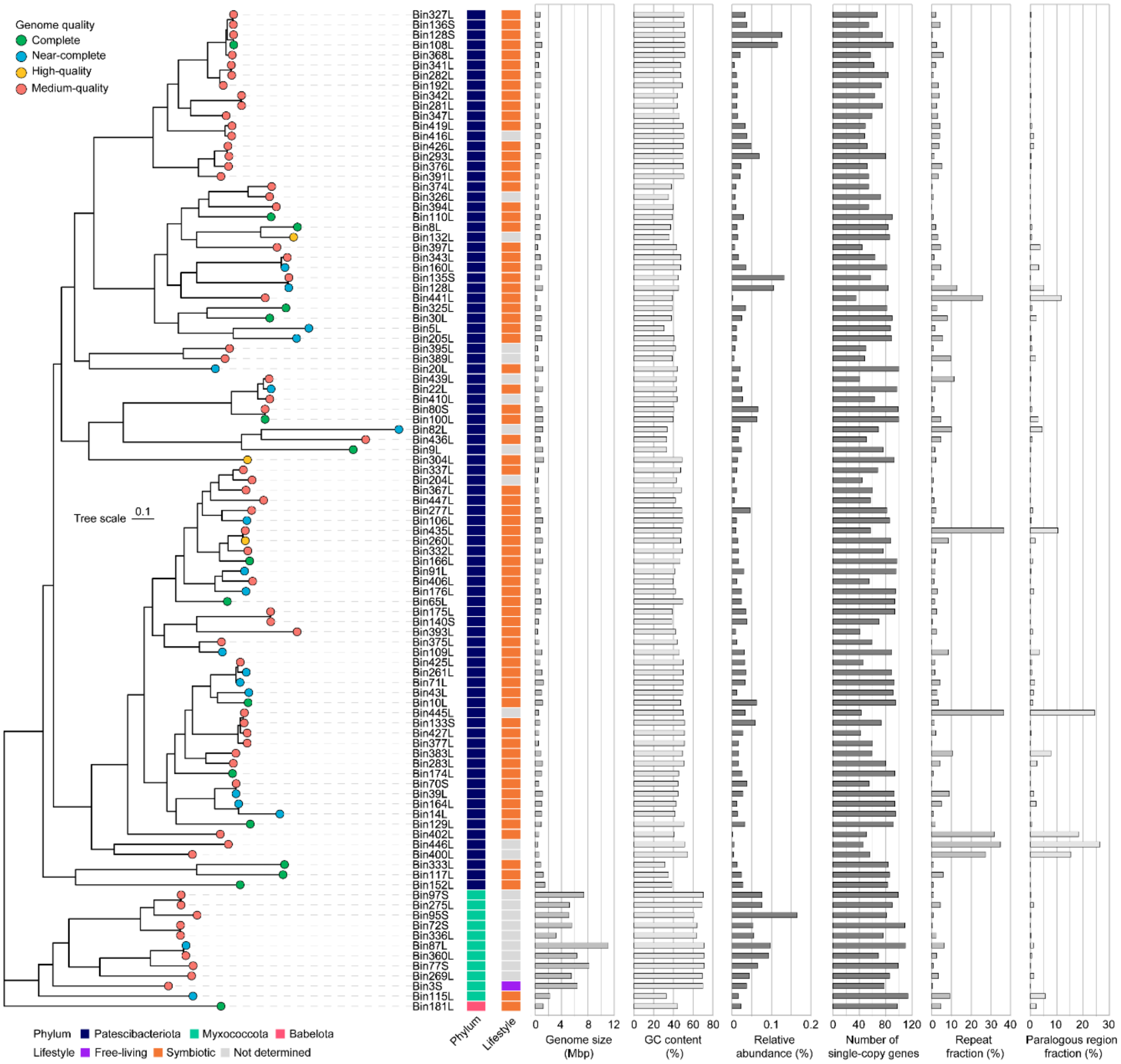

**Supplementary Fig. 1: Phylogenomic diversity and genomic characteristics of Patescibacteriota and Myxococcota MAGs.** Tree tip colors indicate genome quality, and the adjacent strips indicate phylum-level affiliation and lifestyle classification. The adjacent bar plots show genome size, GC content, mean relative abundance, numbers of single-copy genes, and genomic fractions of repeats and paralogous regions. Detailed information is provided in Supplementary Data 1.

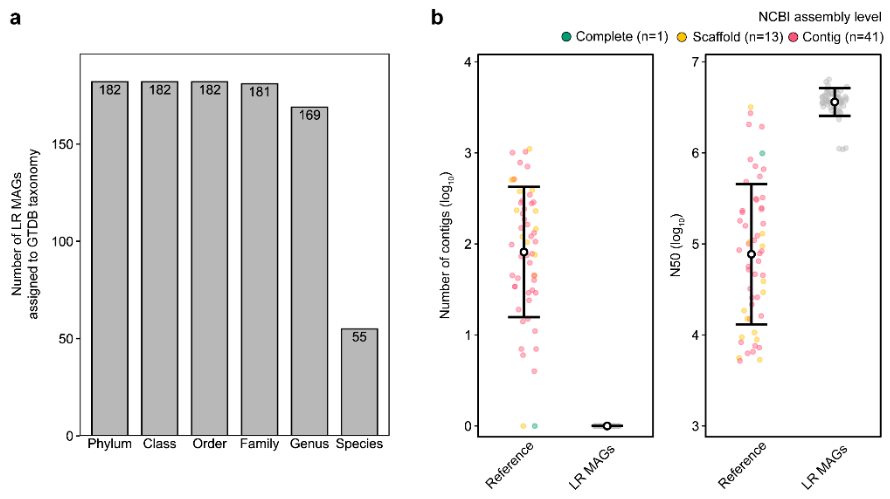

**Supplementary Fig. 2: Taxonomic assignment and reference-genome comparison of (near-)complete LR MAGs. a,** Taxonomic assignment of (near-)complete LR MAGs across GTDB ranks. **b,** Comparison of numbers of contigs and N50 between 55 LR MAGs and their species-matched GTDB reference genomes at different NCBI assembly levels. Detailed information is provided in Supplementary Data 1 and 3.

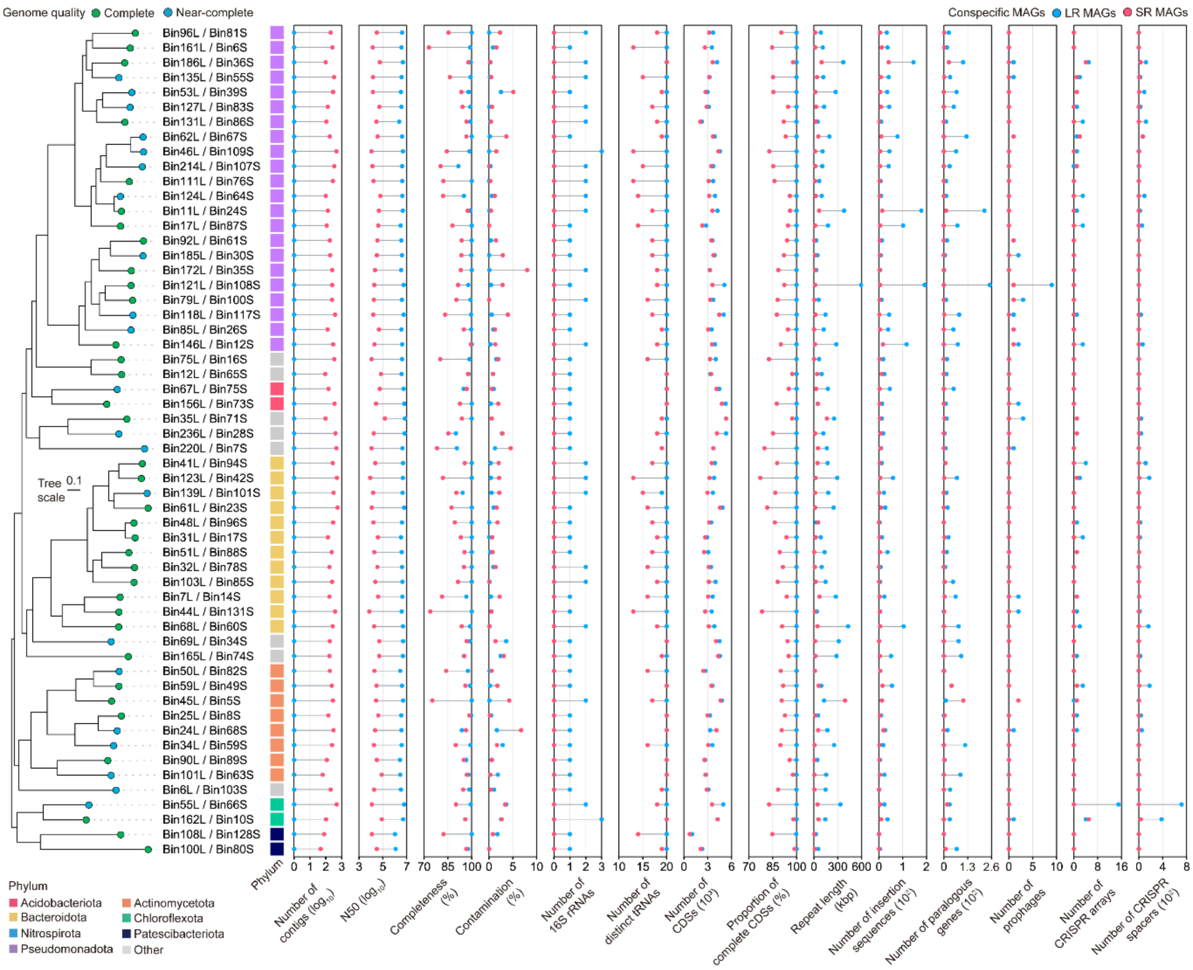

**Supplementary Fig. 3: Phylogenomic tree and genomic comparison of 56 conspecific LR and SR genome pairs.** Tree tip colors indicate LR MAG genome quality. Tip labels indicate paired LR and SR MAG IDs. The adjacent strip indicates phylum-level affiliation. Blue and pink points represent LR and SR MAGs, respectively. Detailed information is provided in Supplementary Data 4.

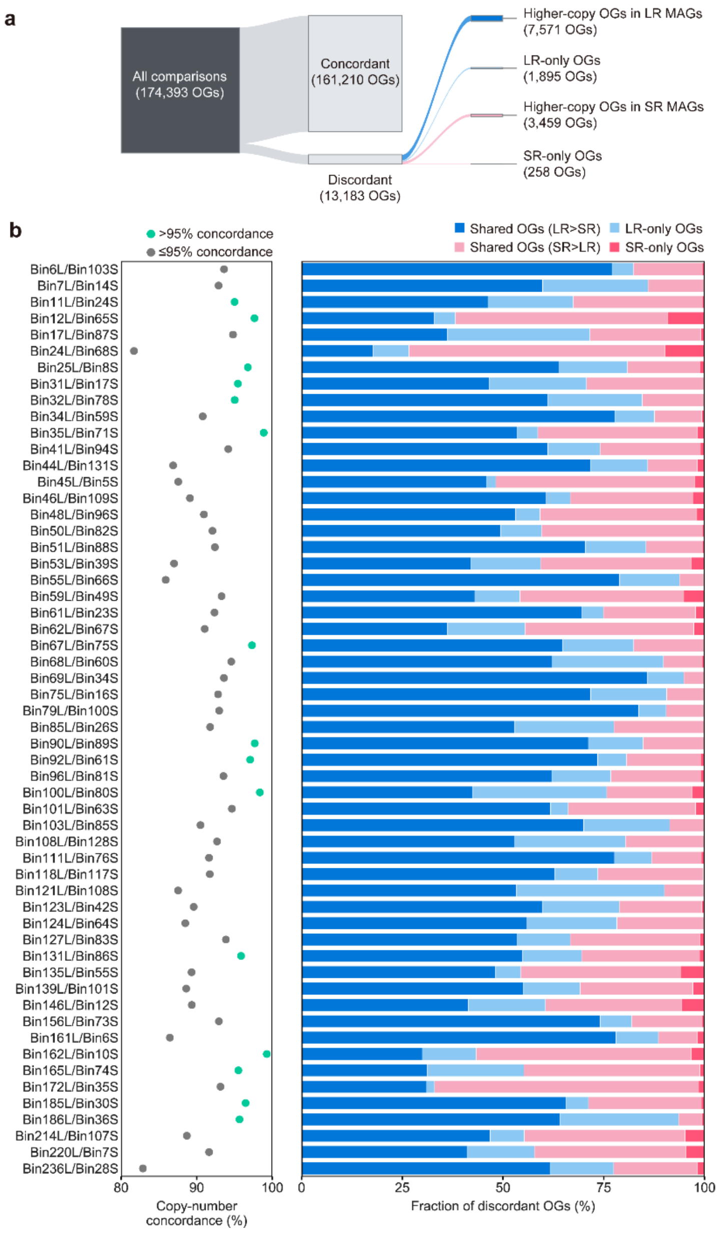

**Supplementary Fig. 4. Gene-family discordance in 56 conspecific genome pairs. a,** Distribution of orthogroup (OG) comparisons according to copy-number concordance and discordance. Discordant OGs were classified as shared OGs with higher copy numbers in LR or SR MAGs, or OGs exclusive to either MAG. Numbers in parentheses indicate OG counts. **b,** Copy-number concordance (left) and proportions of four discordant OG categories (right). Green and grey points indicate >95% and ≤95% concordance, respectively. Detailed information is provided in Supplementary Data 5.

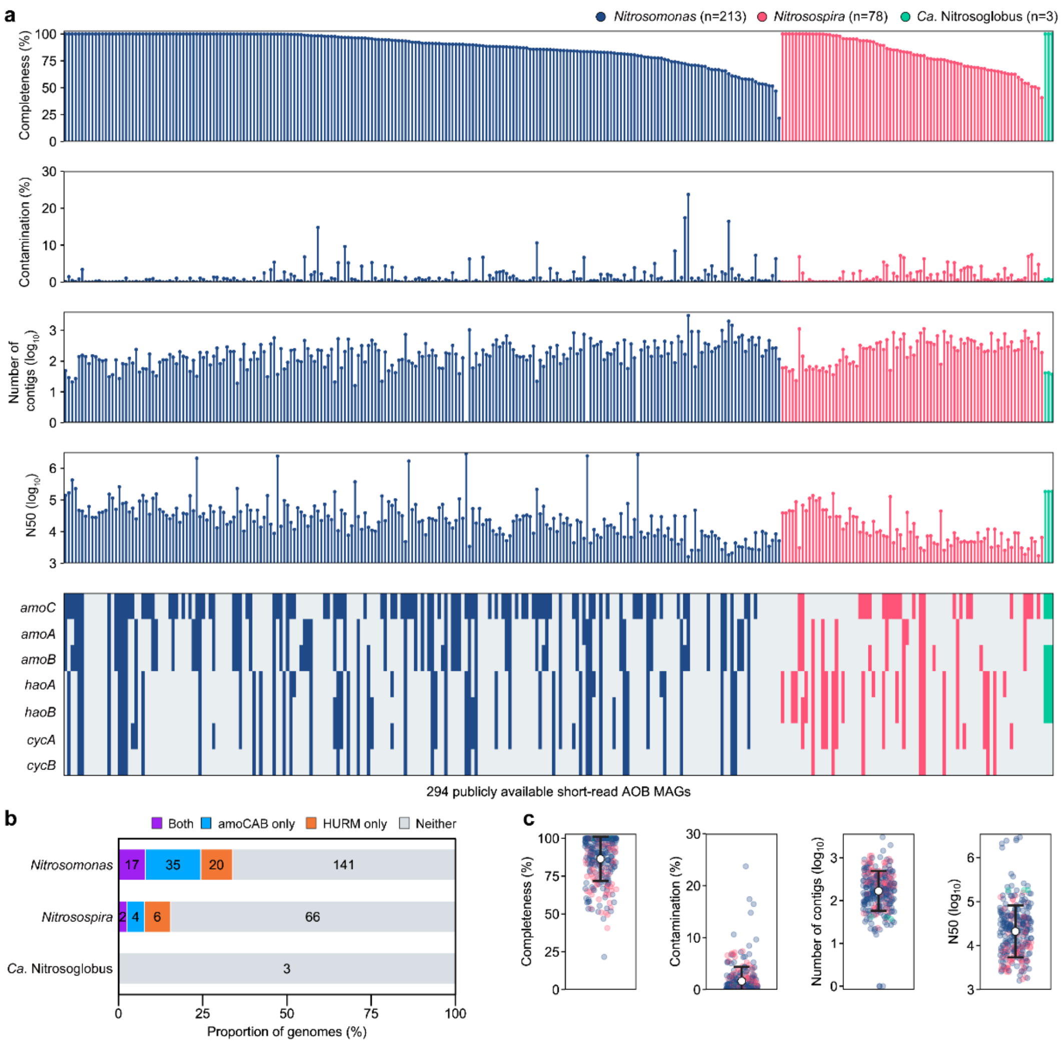

**Supplementary Fig. 5: Genome quality and nitrification marker genes in short-read AOB MAGs. a,** Genome characteristics and nitrification marker genes of 294 publicly available SR AOB MAGs. Genomes are grouped by genus and ordered by decreasing completeness within each genus. Blue, pink and green indicate Nitrosomonas, Nitrosospira and Ca. Nitrosoglobus, respectively. Filled and open cells indicate gene presence and absence, respectively. **b,** Fractions of MAGs encoding both, either or neither of amoCAB and HURM. Numbers in bar indicate genome counts. **c,** Distributions of N50 and numbers of contigs. Grey points indicate individual MAGs, and open black circles with error bars indicate mean±s.d. Detailed information is provided in Supplementary Data 8.

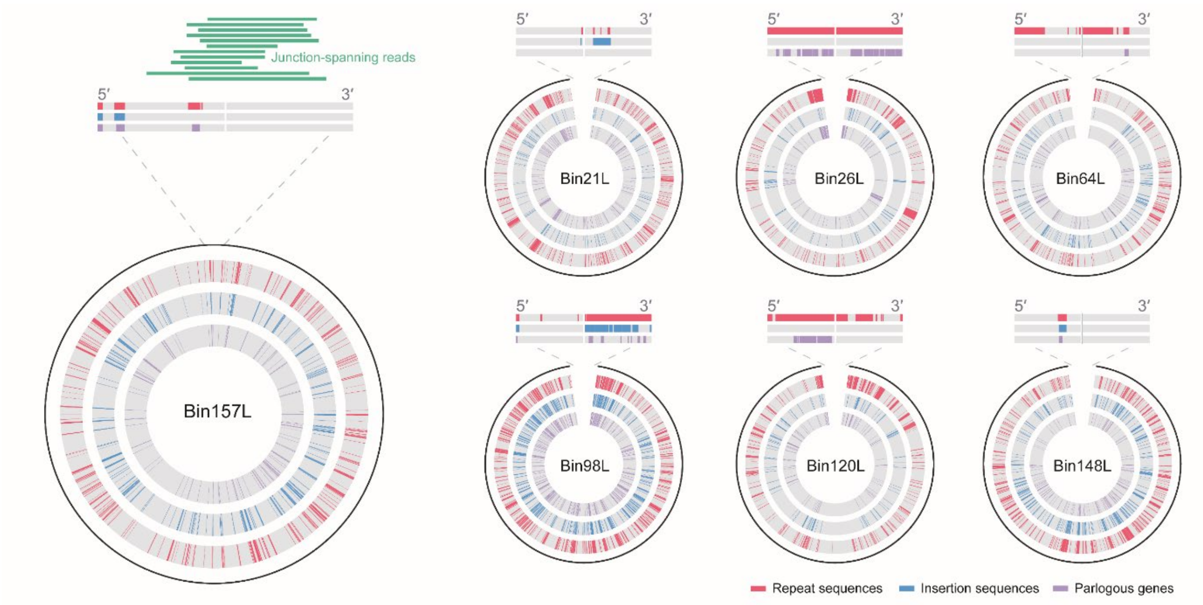

**Supplementary Fig. 6: Validation of genome closure in *Ca.* Accumulibacter bin157L using junction-spanning reads.** Enlarged views of the contig termini in the circular maps of one complete (bin157L) and six near-complete MAGs representing *Ca.* Accumulibacter genus. Only bin157L was reported as circular by the assembler, and this circularity was supported by 12 junction-spanning HiFi reads. The circular maps also show the distributions of repetitive regions, insertion sequences and paralogous genes. Detailed information is provided in Supplementary Data 11.

**Supplementary Table 1. Genome-quality criteria adapted from MIMAG standards.**

| Genome quality | Criteria |
| --- | --- |
| Complete | - Single contig in the absence of ambiguous bases<br>- High-confidence circularity (assembler-reported and HiFi-supported)<br>- Absence of assembly errors identified by HiFi-read validation (phantom sequences, false closure, false duplication, excessive repeats and taxonomic chimeras; Supplementary Note 2)[a]<br>- Presence of the 5S, 16S, and 23S rRNA genes<br>- Presence of at least 18 unique tRNAs<br>- Genome completeness > 90%<br>- Genome contamination < 5% |
| Near-complete | - MAGs satisfying all complete criteria except for high-confidence circularity |
| High-quality | - Multiple contigs, or a single contig with assembly errors<br>- Presence of the 5S, 16S, and 23S rRNA genes<br>- Presence of at least 18 unique tRNAs<br>- Genome completeness > 90%<br>- Genome contamination < 5% |
| Medium-quality | - Genome completeness ≥ 50%<br>- Genome contamination <10%<br>- Not meeting the criteria for higher categories |

[a]High-confidence clipping sites and unsupported variants were not used as classification criteria. Clipping sites were used for classification only when corroborated by cross-contig connections (false closure) or taxonomic transitions (taxonomic chimeras), and unsupported variants were regarded as base-level errors not reflected in genome completeness, contamination or structural integrity.

**Supplementary Table 2. Sequencing and assembly statistics for LR and SR metagenomics.**

| Data type | Platform | Sample / Assembler | Number of sequences ($10^6$) | Total sequence length (Gbp) | Minimum length (bp) | Mean length (bp) | Maximum length (bp) | N50 (bp) | Q30 (%) |
|---|---|---|---|---|---|---|---|---|---|
| Read | LR | 1 | 1.60 | 24.1 | 112 | 15,051 | 59,203 | 15,385 | 94.0 |
| | | 2 | 1.52 | 25.1 | 99 | 16,561 | 63,504 | 17,200 | 93.2 |
| | | 3 | 1.57 | 25.9 | 94 | 16,490 | 63,676 | 17,105 | 93.3 |
| | | 4 | 1.45 | 23.9 | 73 | 16,545 | 66,302 | 17,154 | 93.2 |
| | SR | 1 | 117.88 | 17.8 | 151 | 151 | 151 | 151 | 93.3 |
| | | 2 | 112.92 | 17.1 | 151 | 151 | 151 | 151 | 93.4 |
| | | 3 | 119.33 | 18.0 | 151 | 151 | 151 | 151 | 93.1 |
| | | 4 | 100.65 | 15.2 | 151 | 151 | 151 | 151 | 93.4 |
| Contig | LR | Hifiasm_meta | 0.16 | 11.7 | 7,498 | 75,356 | 8,445,622 | 88,953 | |
| | | metaFlye | 0.13 | 8.5 | 361 | 67,132 | 11,037,590 | 97,476 | |
| | | metaMDBG | 0.34 | 12.0 | 3,296 | 35,442 | 11,102,274 | 50,774 | |
| | SR | MetaSPAdes | 12.62 | 6.9 | 56 | 549 | 611,950 | 771 | |

For each SR sample, the Q30 percentage was calculated as the base-weighted mean of the forward and reverse reads.
